# Cytoplasmic accumulation of a splice variant of hnRNPA2/B1 contributes to FUS-associated toxicity in a mouse model of ALS

**DOI:** 10.1101/2024.10.17.618866

**Authors:** S. Rossi, M. Milani, I. Della Valle, S. Bisegna, V. Durante, M. Addesse, E. D’Avorio, M. Di Salvio, A. Serafino, G. Cestra, S. Apolloni, N. D’Ambrosi, M. Cozzolino

## Abstract

Genetic and experimental findings point to a crucial role of RNA dysfunction in the pathogenesis of Amyotrophic Lateral Sclerosis (ALS). Evidence suggests that mutations in RBPs such as FUS, a gene associated with ALS, affect the regulation of alternative splicing. We have previously shown that the overexpression of wild-type FUS in mice, a condition that induces ALS-like phenotypes, impacts the splicing of hnRNP A2/B1, a protein with key roles in RNA metabolism, suggesting that a pathological connection between FUS and hnRNP A2/B1 might promote FUS-associated toxicity. Here we report that the expression and distribution of different hnRNP A2/B1 splice variants are modified in the affected tissues of mice overexpressing wild-type FUS. Notably, degenerating motor neurons are characterized by the cytoplasmic accumulation of splice variants of hnRNP A2/B1 lacking exon 9 (hnRNP A2b/B1b). *In vitro* studies show that exon 9 skipping affects the nucleocytoplasmic distribution of hnRNP A2/B1, promoting its localization into stress granules (SGs), and demonstrate that cytoplasmic localization is the primary driver of hnRNP A2b recruitment into SGs and cell toxicity. Finally, boosting exon 9 skipping using splicing switching oligonucleotides exacerbates disease phenotypes in wild-type FUS mice. Altogether, these findings reveal that alterations of the nucleocytoplasmic distribution of hnRNP A2/B1, driven by FUS-induced splicing changes, likely contribute to motor neuron degeneration in ALS.

## Introduction

Disturbed nucleocytoplasmic localization of RNA binding proteins (RBPs) stands out as a pivotal aspect in the pathogenesis of Amyotrophic Lateral Sclerosis (ALS), a progressive neurodegenerative condition marked by the degeneration of motor neurons, resulting in gradual muscle wasting and weakness ^1–4^. A notable example is TDP-43, a member of the hnRNP family of RBPs, which undergoes nuclear-to-cytoplasmic translocation in most familial and sporadic ALS cases ^5^. This leads to the loss of nuclear splicing regulation of TDP-43 target sequences, including intronic sequences incorrectly incorporated into mature mRNA as “cryptic exons”. This mechanism generates dysfunctional proteins critical for neuronal activity, thus contributing to disease pathogenesis ^6^. In addition, cytoplasmic-accumulated TDP-43 localizes into stress granules (SGs), RNA-protein condensates forming in the cytoplasm under stress stimuli. In ALS conditions, these granules exhibit impaired dynamics, and act as seeds for the accumulation and aggregation of several other RBPs, thus inducing a cascade of alterations in the regulatory network of protein and RNA metabolism homeostasis ^7–9^.

The gain of cytoplasmic activities and the loss of nuclear functions may also explain the toxicity of ALS-associated FUS, an hnRNP responsible for about 5% of genetic ALS cases ^10–13^. Indeed, mutations in the nuclear localization signal (NLS) or in the prion-like domain (PrLD) result in an increased cytoplasmic localization of FUS ^12^. This promotes the recruitment of FUS into SGs, potentially evolving into pathological inclusions found in post-mortem brain and spinal cord tissues from patients harbouring FUS mutations ^8,14^. Moreover, upregulation of FUS wild-type protein expression due to mutations in its 3′ UTR regulatory region causes ALS in patients and ALS-like disease in animal models ^15–18^, further supporting a gain-of-function mechanism of neurotoxicity. However, loss-of-function mechanisms are likely involved, as suggested by the alterations in the splicing of several target genes identified in cellular and animal models of FUS-ALS ^19–21^. Interestingly, the genes that are affected in their splicing and/or expression upon FUS mutation are enriched in functional categories related to RNA metabolism ^21,22^, implying that pathological FUS may affect the overall expression, distribution, and activity of other RBPs, that might in turn contribute to motor neuron degeneration.

By employing cultured cells and transgenic mice overexpressing human wild-type FUS (hFUS), here we uncover the existence of a functional connection between FUS and hnRNP A2/B1, an RBP involved in various steps of RNA metabolism, including alternative splicing. In hFUS mice, disease progression is marked by the accumulation of hnRNP A2/B1 splicing isoforms lacking exon 9 (hnRNP A2b/B1b), which preferentially localize in the cytoplasm of degenerating motor neurons. *In vitro* experiments demonstrate that the hnRNP A2b variant exhibits an increased propensity to relocalize into the cytoplasm, and that its de-localization is sufficient to drive SGs formation and cell toxicity. Finally, disease phenotypes in hFUS mice worsen upon treatment with splicing switching oligonucleotides that enhance exon 9 skipping. Overall, these findings support the existence of a pathological cascade orchestrated by FUS and hnRNP A2/B1 and strengthen the idea that the functional network connecting RBPs is widely affected by ALS conditions.

## Results

### hnRNP A2/B1 splicing isoforms are modulated in the spinal cord of symptomatic hFUS mice

We have previously found that ALS disease in mice overexpressing wild-type human FUS (hFUS mice) is characterized by a rearrangement in the expression of hnRNP A2/B1 RNA splicing isoforms ^20^. To investigate the relevance of this process, we analyzed the splicing of hnRNP A2/B1 at different time points of disease progression in hFUS mice. As shown in Figure 1A, B, the spinal cord from hFUS transgenic mice at the end stage of the disease (39-41 days) is marked by a significant increase in the expression of hnRNP A2/B1 isoforms lacking exon 9 (A2b and B1b), with concomitant downregulation of exon 9-containing isoforms (A2 and B1). The skipping of hnRNP A2/B1 exon 9 occurs concomitantly with the appearance of disease symptoms in hFUS mice (31-36 days) and is sustained until the end-stage phase of the disease. No changes in the alternative splicing of exon 9 are evident at the pre-symptomatic phase of the disease, suggesting that this event is linked to the disease onset and progression. Importantly, the alternative splicing of exon 2, which underlies the expression of B1 and B1b isoforms, remains unchanged throughout the disease course, indicating that exon 9 splicing is specifically affected.

**Figure 1.**
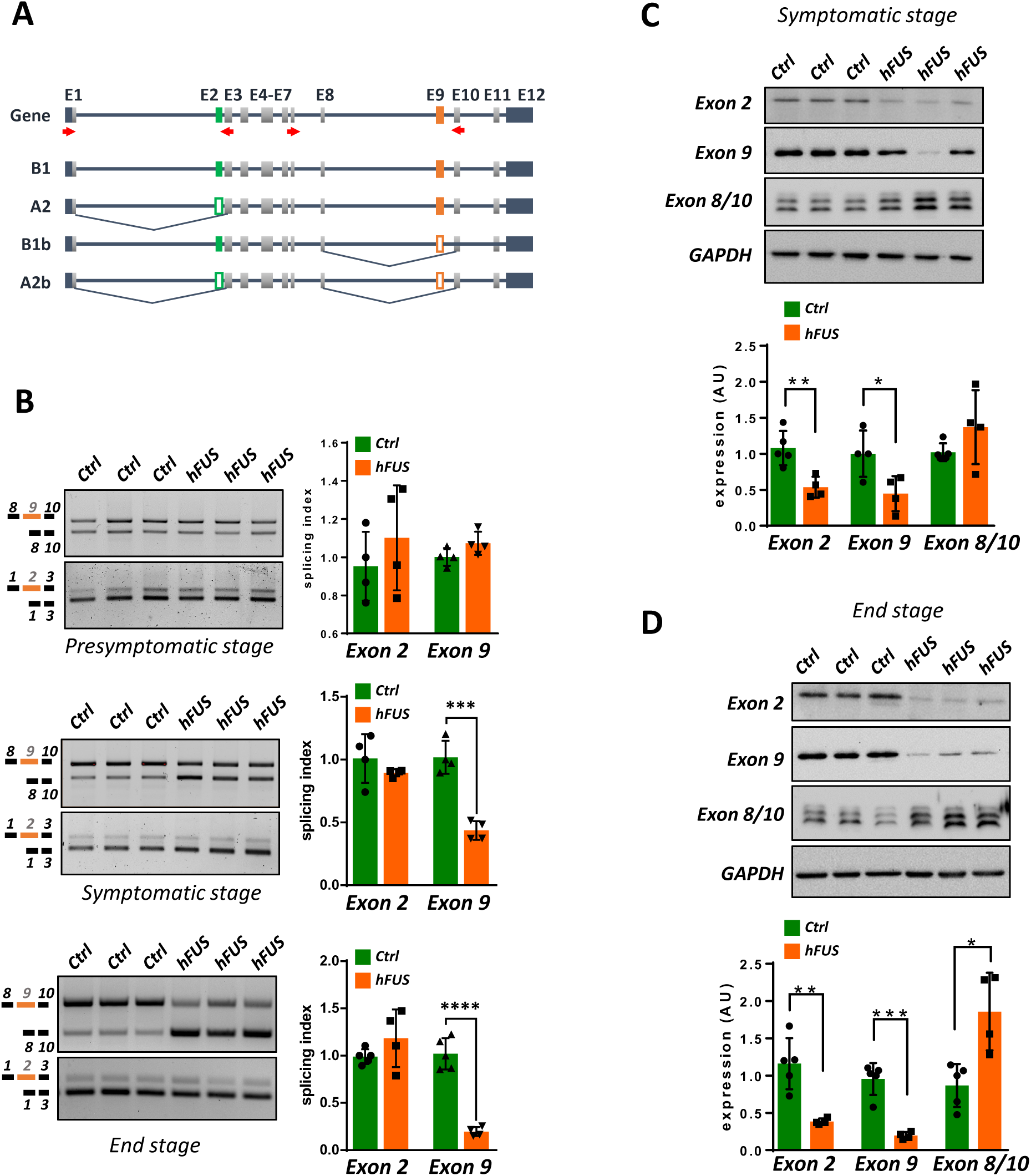
hnRNP A2/B1 isoforms are modulated along with symptoms onset in the spinal cord of diseased hFUS mice. (A) Schematic representation of alternative splicing of exon 2 (green) and exon 9 (orange) of hnRNP A2/B1. The filled rectangles represent included exons, while the empty rectangles represent skipped exons. Arrows represent the specific primers used for cDNA amplification. (B) The alternative splicing pattern of hnRNP A2/B1 exon 2 and exon 9 was monitored by semiquantitative RT-PCR analysis in spinal cords from hFUS transgenic mice, along with age-matched non-transgenic (Ctrl) animals, at the pre-symptomatic, symptomatic and end stages of the disease. Bands were quantified through densitometric analysis, and a splicing index was calculated as the ratio between the upper and lower band and plotted considering the corresponding ratio in a Ctrl mouse equal to 1. Data are expressed as means ± SD (n=4-5 mice/group). Statistical significance was calculated by Student’s t-test, ***p<0.001, ****p<0.0001. (C-D) Lumbar spinal cord lysates from control (Ctrl) and hFUS mice at the symptomatic (C), and end-stage (D) phases of the disease were subjected to western blot analysis using anti-exon 2, anti-exon 9, and anti-exon 8/10 antibodies. GAPDH was used as a loading control. Data are expressed as means ± SD (n=4/5 mice/group) considering the relative expression of a Ctrl mice equal to 1. Statistical significance was calculated by Student’s t-test referred to Ctrl, *p<0.05, **p<0.01, ***p<0.001.

To verify if these splicing changes modify the overall expression of hnRNP A2/B1 proteins and/or the expression of specific protein isoforms, we made use of antibodies that can distinguish the different isoforms (Supplementary Figure 1). As shown in Figure 1C, at a symptomatic phase of the disease the expression levels of hnRNP A2/B1 isoforms containing exon 2 and exon 9 are significantly decreased in hFUS mice compared to control, non-transgenic animals, as assessed by an anti-exon 9 antibody, recognizing B1 and A2 isoforms, and an anti-exon 2 antibody, which binds B1 and B1b isoforms. On the contrary, exon-9-devoid isoforms, recognized by an antibody raised against a fusion epitope between exon 8 and 10 (anti-exon 8/10 antibody), show an increasing expression trend. These results were confirmed when the same analysis was performed at the terminal phase of the disease (Figure 1D), where anti-exon2, and anti-exon 9-immunoreactive isoforms are strongly downregulated, while anti-exon 8/10 signal is significantly upregulated in hFUS mice compared to control animals. Notably, these effects appear enhanced at this stage of the disease, suggesting that the observed alterations in the expression of hnRNP A2/B1 match disease progression. Overall, these results show that ALS disease in hFUS mice is characterized by a shift in the expression of hnRNP A2/B1 towards isoforms lacking exon 9 (either A2b, B1b, or both), consistent with the splicing data.

### Degenerating spinal cord MNs display cytoplasmic accumulation of A2b/B1b

To provide a detailed characterization of the expression of hnRNP A2/B1 isoforms in different cell types, we performed confocal immunofluorescence analysis on spinal cord sections of both non-transgenic and end-stage hFUS mice. As reported in Figure 2A, the anti-exon 9 antibody shows nuclear staining in both grey and white matter of the spinal cord ventral horn from control animals, with SMI32-positive motor neurons clearly expressing hnRNP A2/B1 isoforms containing exon 9. Notably, a clear decrease in exon 9 signal is observed in hFUS mice. In addition, several GFAP-positive astrocytes display anti-exon 9 immunoreactivity, as well as few CD11b-positive microglia/macrophages (Figure 2A), indicating that exon 9-containing variants are present in glial cells. The immunostaining of exon2-containing isoforms (B1 and B1b) gave similar results (Fig. 2B). Indeed, almost all SMI32-positive motor neurons in non-transgenic, and the remaining motor neurons in hFUS mice express exon 2-containing isoforms, while only a few GFAP-positive astrocytes and CD11b-positive microglia show immunoreactivity for these isoforms. As expected, hFUS mice display a remarkable decrease in B1/B1b staining, in accordance with western blot results. Finally, immunostaining of spinal cord sections with the anti-exon 8/10 antibody reveals a clear signal in SMI-32-positive neurons in the grey matter of hFUS mice as well as into ctrl, non-transgenic mice (Figure 2C). Most importantly, in SMI32-positive neuronal cells, A2b/B1b isoforms are affected in their distribution, ranging from a preferential nuclear localization in neurons of control mice to a clear cytoplasmic localization in neurons from hFUS mice (Figure 2C). This is particularly evident in the soma of hFUS motor neurons, where the cytoplasmic staining of hnRNP A2/B1 is condensed into granules that might be reminiscent of condensed/insoluble species (Fig. 2C). Finally, in the white matter of hFUS mice, GFAP- and CD11b-positive cells display a clear exon 8/10 signal, suggesting that glial cells might contribute to the increased expression of A2b/B1b isoforms during disease progression (Fig 2D). Taken together, results from these experiments show that hnRNP A2/B1 isoforms have a widespread distribution in the spinal cord cell populations, with exon 9-containing isoforms mostly expressed in neuronal cells, while exon 9-devoid isoforms also expressed in reactive glial cells. Above all, these data show that the pathological process is marked by a significant rearrangement of hnRNP A2/B1 isoform expression and distribution in the spinal cord during disease progression, further suggesting that a functional impairment of hnRNP A2/B1 might have a role in this process.

**Figure 2.**
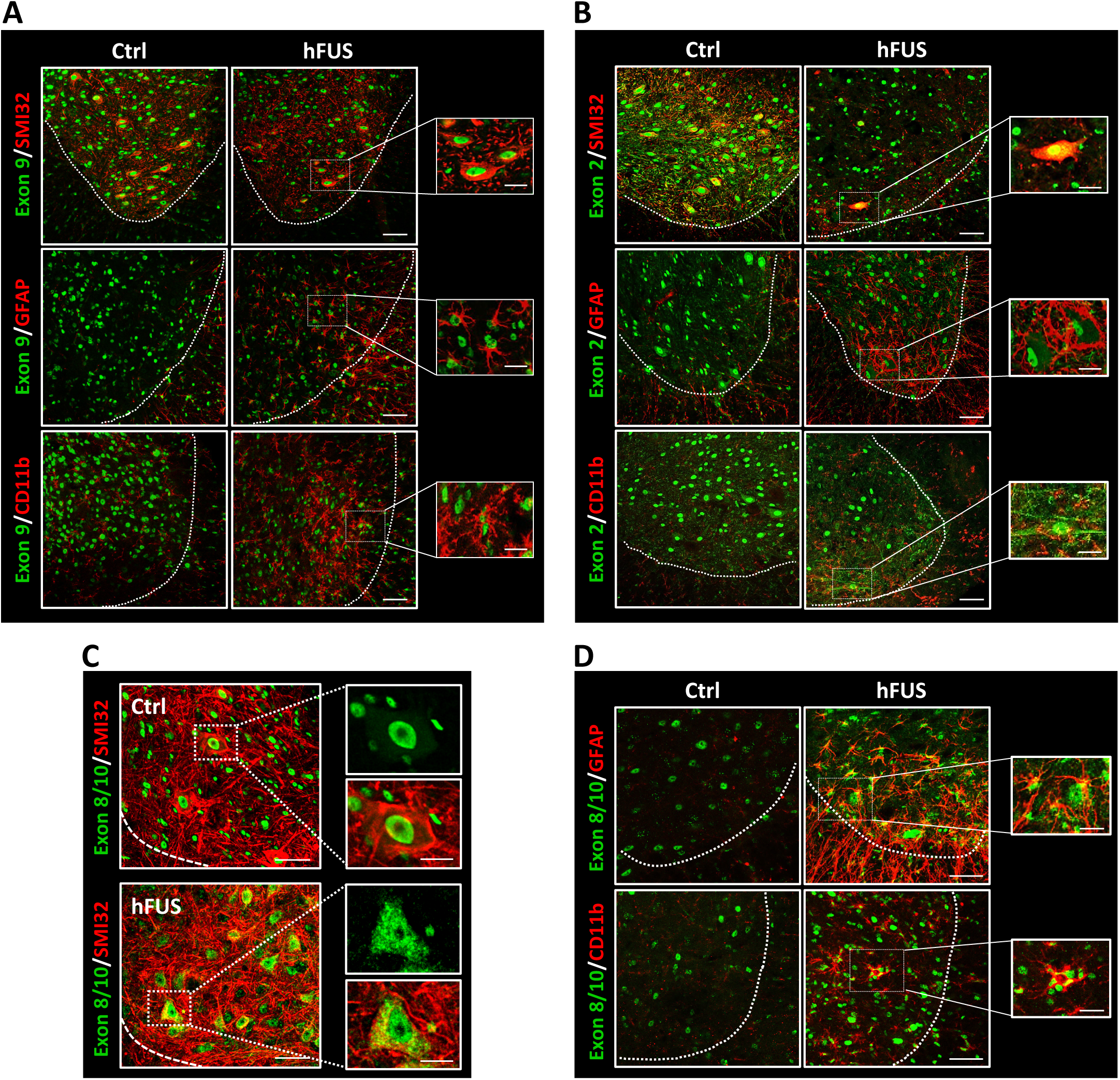
Exon 9 and exon 2 containing isoforms are present on neuronal and non-neuronal cells in the spinal cord of FUS mice. (A, B, C, D) Immunofluorescence staining on spinal cord sections of non-transgenic (Ctrl) and end stage hFUS mice (n=4 mice/group) with an antibody against Exon 9 (green) (A), Exon 2 (green) (B) or Exon 8/10 (green) (C, D) and SMI32 (red), GFAP (red) and CD11b (red). The dotted white lines mark the separation between white and grey matter of the spinal cord. Magnifications of the highlighted areas are also shown. Scale bar: 50 µm, 20 µm (inset).

### Exon 9 skipping causes diffuse cytoplasmic distribution of hnRNP A2/B1

Previous experiments suggest that exon 9 skipping from hnRNP A2/B1 promotes cytoplasmic localization. To verify this possibility, we analyzed the subcellular distribution of the four hnRNP A2/B1 isoforms when expressed in cultured cells. HeLa cells were transfected with HA-tagged hnRNP A2/B1 isoform constructs, and analyzed by immunofluorescence using an anti-HA antibody, which detects exclusively exogenous hnRNP A2/B1 isoforms, and an antibody anti-TIA1, which is commonly used as a marker of SGs. Exon 9-containing isoforms (A2 and B1) have a marked nuclear localization (Figure 3 A,B), while isoforms lacking exon 9 (A2b and B1b) additionally localize in the cytoplasm of transfected cells. Interestingly, a significant fraction of cells transfected with A2b isoform and, to a lesser extent, B1b isoform, displays cytoplasmic SGs that are positive to HA staining, indicating that cytoplasmic hnRNP A2/B1 accumulates into SGs (Figure 3A,B). Consistently with immunofluorescence analysis, nucleo-cytoplasmic fractionation shows that both A2 and A2b isoforms are expressed in the nuclear fraction, while A2b is clearly detectable also in the cytoplasmic fraction, differently from A2 (Figure 3C). Similarly, B1 and B1b are mostly present in the nuclear fractions, with B1b showing a slight increase in the cytoplasm compared to B1 (Figure 3C). These results show that the lack of exon 9 promotes cytoplasmic localization of hnRNP A2/B1 in HeLa cells. A similar outcome was obtained when mouse primary cortical neurons were subjected to analogous experiments. Indeed, exon 9-lacking isoforms localize into the soma and the neurites of transfected neurons, while exon 9-containing isoforms are mostly nuclear (Supplementary Figure 2).

**Figure 3.**
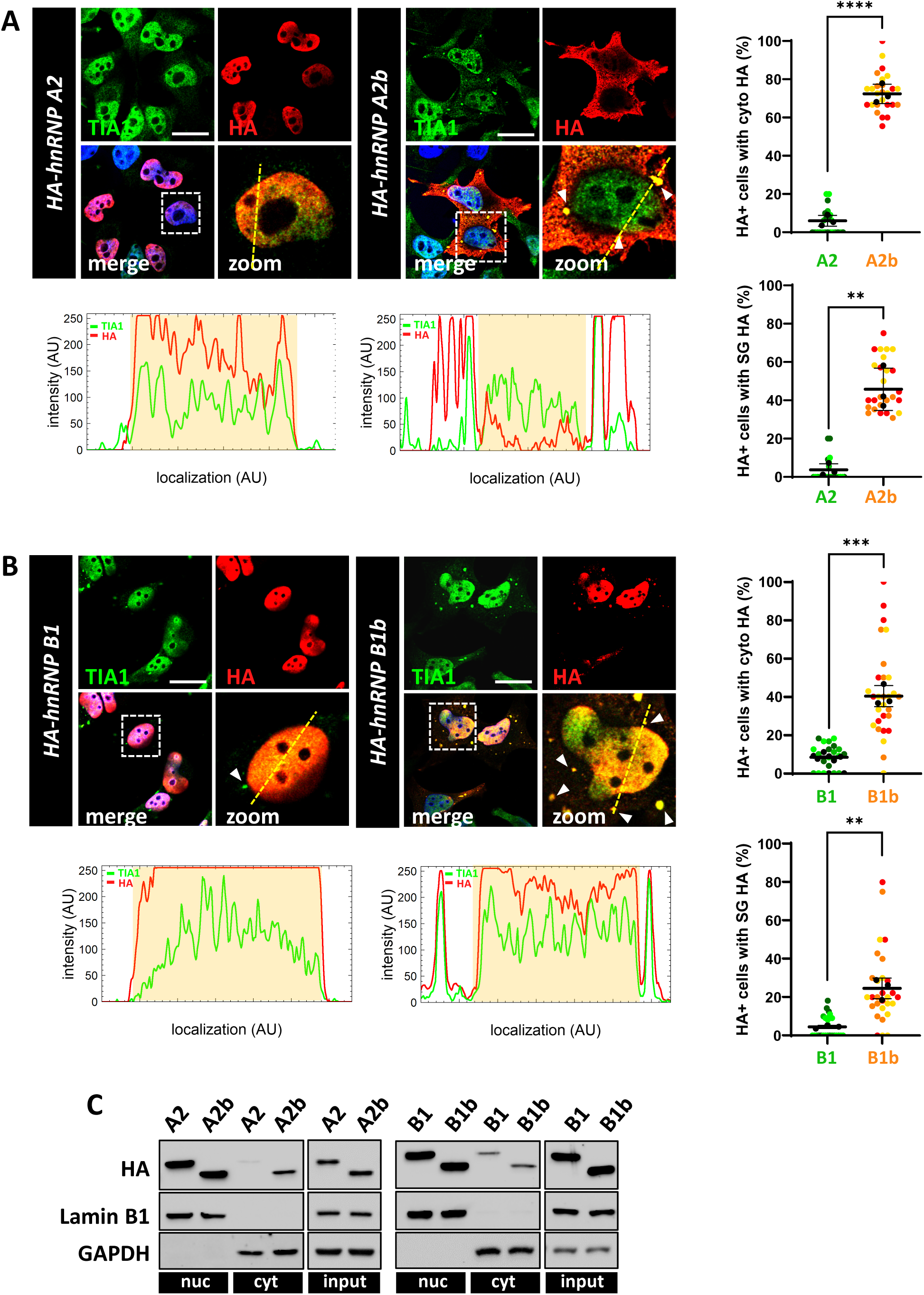
Isoforms containing exon 9 display nuclear localization, while isoforms lacking exon 9 localize in cytoplasm and/or stress granules in cultured cells. HeLa cells were transfected with the HA-tagged hnRNP A2, hnRNP A2b (A), B1 and B1b (B) isoform constructs and analysed 24h after transfection by immunofluorescence, using an anti-HA antibody (red) and anti-TIA1 antibody (green). White arrowheads point at SGs. Magnifications of the highlighted areas are also shown (zoom). Scale bar: 20 μm. For fluorescent distribution across the cell, ImageJ (NIH) was used. A straight line was overlaid across the cell and then the fluorescent intensity was measured across the line using the built-in function. Graphs (lower panels) represent fluorescent intensity across the line in images; the yellow shaded area denotes the nucleus. SuperPlots on the right show the percentage of cells with cytosolic HA signal or cells where the HA signal colocalizes with TIA1-positive stress granules, calculated for both the A2/A2b (A) and B1/B1b isoforms (B). The distribution of measures from n=3 independent experiments is reported, with each biological replicate color-coded: the mean value from each of the three replicates is represented by black dots, and the mean ± SD of the three replicates is shown as a black line. Statistical significance was calculated by Student’s t-test, **p<0.01, ***p<0.001, ****p<0.0001. (C) Total protein extracts (input), as well as the nuclear (nuc) and cytosolic (cyt) fractions from HeLa cells transfected with HA-tagged hnRNP A2/B1 isoform constructs were analyzed by western blot. hnRNP A2/A2b/B1/B1b expression levels were monitored using an anti-HA antibody. GAPDH, lamin B1 and β-actin levels were included to assess the purity of the cytosolic, nuclear, and total fractions.

### Exon 9 skipping increases hnRNP A2/B1 recruitment to stress granules

HnRNP A2/B1 has been shown to localize into cytoplasmic SGs under stress conditions, like those induced by sodium arsenite (NaAs) treatment ^23,24^. Further, in the previous experiments we have highlighted the ability of exon 9-lacking isoforms to accumulate into SGs at a steady state in a significant fraction of cells. Therefore, to evaluate whether the presence or absence of exon 9 and/or exon 2 might influence their propensity to be recruited into SGs, we analyzed their behaviour under stress conditions. As shown in Figure 4, A2 isoform localizes into the cytoplasm and accumulates into SGs in a time-dependent manner under NaAs treatment. Interestingly, this tendency is dramatically enhanced for the A2b isoform, which shows increased stress granule localization as early as 20’ treatment with NaAs (Figure 4 A,B). Further, while B1 isoform does not localize in the cytoplasm nor into SGs even after 60’ treatment with NaAs, B1b tends to re-localize into SGs in a time-dependent manner comparably to A2b, although to a lesser extent (Supplementary Figure 3). Similar results were obtained in neuronal SH-SY5Y cells (Supplementary Figure 4). Altogether, these results indicate that hnRNP A2/B1 localization into cytoplasmic SGs is antagonized by the presence of exon 9. The distribution of hnRNP A2/B1 isoforms was confirmed by nucleo-cytoplasm fractionation. A2 and A2b isoforms were chosen as they show more evident differences in their subcellular localization. Consistently with immunofluorescence analysis, both A2 and A2b demonstrate a progressive increase in their expression in the cytoplasmic fraction under NaAs treatment (Figure 4C). However, the levels of A2b in the cytoplasmic fraction are higher compared with those of A2 (Figure 4C).

**Figure 4.**
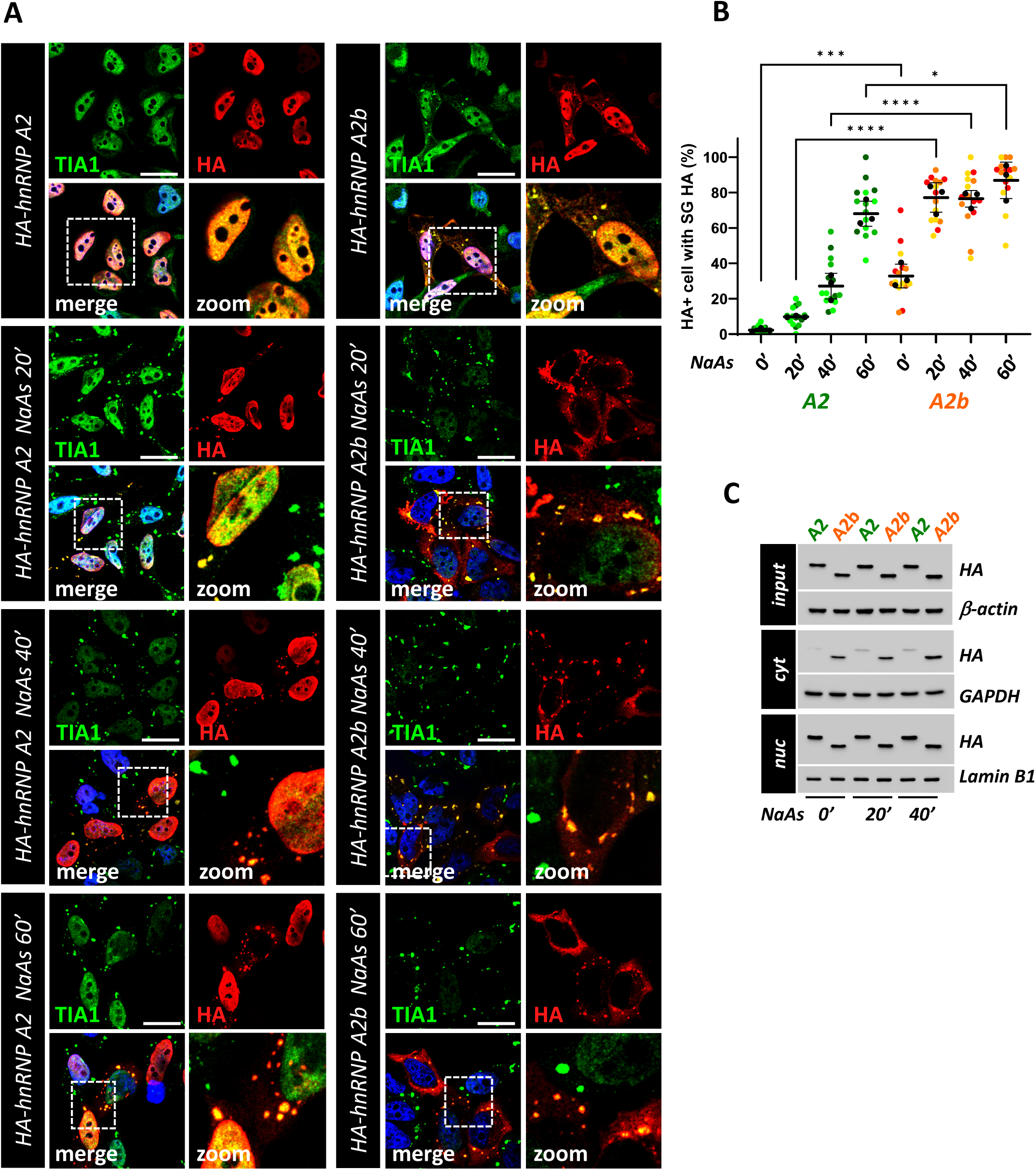
A2b shows increased localization in stress granules compared to A2 under NaAs treatment. (A) HeLa cells were transfected with the HA-tagged hnRNP A2 and hnRNP A2b isoform constructs for 24h, untreated or treated with 0.5 mM sodium arsenite (NaAs) for 20, 40 and 60 minutes and analysed by immunofluorescence using an anti-HA antibody (red) and anti-TIA1 antibody (green). Nuclei were detected by DAPI staining (blue). Scale bar: 20 μm. Magnifications of the highlighted areas are also shown (zoom). (B) SuperPlots showing the percentage of cells with the HA signal colocalizing with TIA1-positive stress granules. The distribution of measures from n=3 independent experiments is reported, with each biological replicate color-coded: the mean value from each of the three replicates is represented by black dots, and the mean ± SD of the three replicates is shown as a black line. Statistical significance was calculated by One-way ANOVA test, *p<0.05, ****p<0.0001, and the significant differences between A2 and A2b isoforms at the same time point are shown. (C) Total protein extracts (input), as well as the nuclear (nuc) and cytosolic (cyt) fractions from HeLa cells transfected with HA-tagged hnRNP A2 and A2b isoforms constructs, were analysed by western blot. hnRNP A2 and A2b expression levels were monitored using an anti-HA antibody. GAPDH, Lamin B1 and β-actin levels were analyzed to assess for the purity of the cytosolic, nuclear and total fractions.

A point mutation in the hnRNP A2/B1 gene (D290V) has been linked to rare familial ALS cases, and it is known that this mutation promotes the accumulation of hnRNP A2 into SGs ^23^. The D290V mutation localizes near exon 9, into the protein prion-like domain, which is a key determinant for the assembly of hnRNP A2/B1 into ribonucleoprotein granules, including SGs ^23^. For this reason, we compared the behaviour of A2 and A2b in the absence or presence of D290V mutation under stress conditions. As shown in Supplementary Figure 5, at basal conditions no major differences in stress granule localization between wild-type and D290V isoforms were detected. However, after 40’ treatment with NaAs, A2-D290V shows an increased localization into SGs compared to wild-type A2, as already reported ^23^; (Supplementary Figure 5 A,B). Notably, the A2b isoform displays a higher propensity to localize into SGs compared to A2-D290V (Supplementary Figure 5 A,B), and the presence of the D290V mutation does not further enhance this effect.

### Cytoplasmic localization is the driver of hnRNP A2 recruitment into SGs and cell toxicity

Previous results prompted us to verify if the increased localization of the A2b variant into SGs might reflect changes in its propensity to localize into cellular condensates and/or in its solubility, like what has been described for the D290V A2 mutant. As shown in Figure 5 A,B, a higher proportion of detergent-soluble A2b isoform compared to A2 was detected in A2b and A2 transfected cells, respectively. This effect is not antagonized by the D290V mutation, which is indeed able to induce a slight decrease in A2 solubility, but that has no effect on the solubility of the A2b variant.

**Figure 5.**
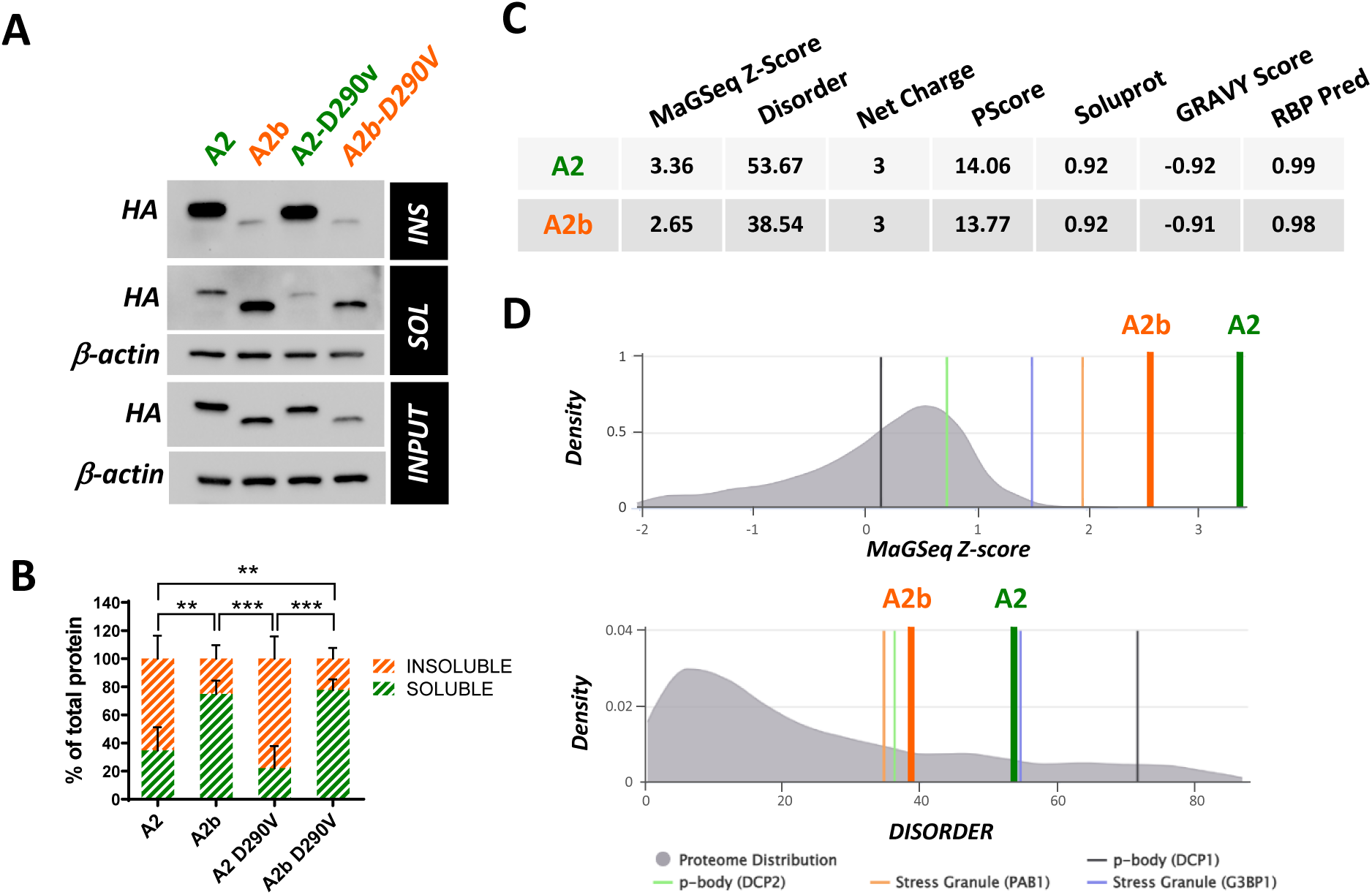
Isoform lacking exon 9 shows increased solubility. Representative western blot (A) and quantification (B) of total (INPUT) protein extracts, as well as the insoluble (INS) and soluble (SOL) fractions from HeLa cells transfected with HA-tagged hnRNP A2, A2b, A2-D290V and A2b-D290V isoforms constructs. Isoform expression levels were monitored using an anti-HA antibody. β-actin levels were used as a loading control. Data are reported as mean value ± SD (n=3 independent experiments). Statistical significance was calculated by Two-way ANOVA, comparing soluble and insoluble fractions between groups. **p<0.01, ***p<0.001. (C) Numerical output scores resulting from GraPES analysis of hnRNP A2 compared to hnRNP A2b, including: MaGSeq Z-score, with higher values indicating increased likelihood of the protein being localized in a biological condensate (a value greater than 0.90 suggests a high propensity); Disorder, representing the percentage of protein residues predicted to be disordered by DISOPRED3; Net charge, that is the overall sum of the positively and negatively charged residues at neutral pH; PScore, reflecting the quantity of π-π interactions, that are linked to the propensity of the protein to phase separate in vitro; Soluprot, a protein solubility score where higher values correspond to higher solubility; GRAVY Score, a measure of protein hydrophobicity; and RBP Pred, a likelihood prediction for a protein to exhibit RNA-binding capabilities. (D) Graphical plots generated by GraPES analysis showing MaGSeq Z-score (upper panel) and Disorder score (lower panel) of hnRNP A2 and hnRNP A2b along the precomputed score distributions of human proteome (shown in gray). The y-axis represents the percentage (density) of proteins associated with a given score (the x-axis). Scores relative to known markers for SGs and processing bodies (p-bodies) are also shown as references.

The solubility of RBPs is often connected to the presence of intrinsically disordered regions (IDRs) which underlie their ability to phase separate into membraneless organelles, including SGs ^25^. We therefore subjected hnRNP A2 splicing isoforms to Granule Protein Enrichment Server (Grapes) analysis, a prediction tool that calculates propensity scores for protein localization into cellular condensates ^26^. As shown in Figure 5, A2 ranks among the proteins with the highest degree of disorder and tendency to localize into SGs, in agreement with previous experimental evidence ^23^. Importantly, removing exon 9 clearly reduces protein disorder and decreases A2 propensity to stress granule localization. Overall, these data indicate that the increased ability of A2b to localize into SGs is not strictly related to an altered solubility and/or disorder caused by the absence of exon 9 and suggest that nucleo-cytoplasmic localization might be directly involved. To verify this possibility, we used A2 and A2b isoforms bearing or lacking exon 11 (Fig. 6A). It has been recently shown that the removal of this exon from hnRNP A2/B1, and the consequent deletion of a portion of the C-terminal M9-nuclear localization signal (NLS) is sufficient to promote cytoplasmic accumulation ^27^. Figure 6B, C shows that, differently from A2, the A2 variant lacking the M9-NLS displays a clear cytoplasmic localization, and is significantly recruited into SGs, similarly to the A2 variant lacking exon 9. On the contrary, an A2b variant that is forced to localize in the nucleus due to the presence of an extra NLS does not localize into SGs. This suggests that exon 9 contributes to the nuclear-cytoplasmic distribution of hnRNP A2/B1 and indicates that the increased cytoplasmic localization due to exon 9 skipping is a major driver of SG localization. Finally, to investigate the functional consequence of cytoplasmic localization, A2 and A2b splicing variants were expressed in SH-SY5Y neuronal cells. As shown in Figure 6D,E, the expression of A2b induces an apoptotic phenotype, as indicated by the increased expression of active caspase 3 and a cleaved form of PARP1, compared to A2 or mock-transfected cells. Importantly, the effect of A2b is mimicked by the A2_ΔNLS variant, indicating that cytoplasmic localization is sufficient to induce cell toxicity. As expected, A2b_ΔNLS, induces apoptosis.

**Figure 6.**
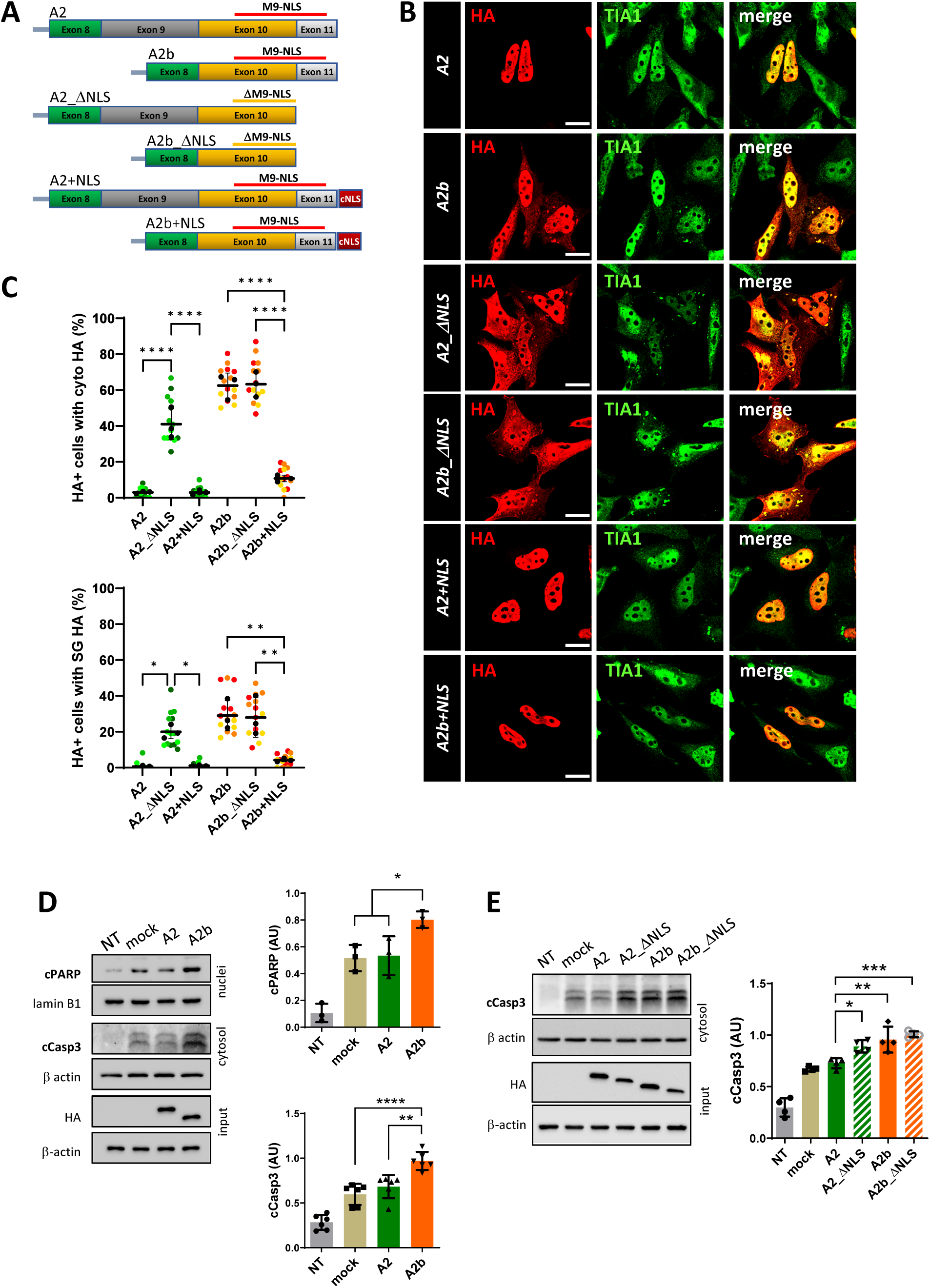
Cytoplasmic delocalization is sufficient to promote stress granule accumulation of hnRNP A2/B1 and cell toxicity. (A) Schematic representation of hnRNP A2/B1 variants used. hnRNP A2 isoforms, with (A2) or without (A2b) exon 9, were further modified to exclude exon 11, which encodes a portion of the nuclear localization signal (M9-NLS) (A2_ΔNLS and A2b_ΔNLS). hnRNP A2 and A2b isoforms fused at the C-terminal with an extra nuclear localization signal (NLS) were also produced. All six variants contain an HA epitope at the N-terminal. (B) Hela cells were transfected with the indicated plasmids and analyzed by immunofluorescence using anti-HA (red) and anti-TIA1 (green) antibodies. A merge of the two signals is also shown. (C) SuperPlots showing the percentage of cells with cytosolic HA signal (upper panel) and cells where the HA signal colocalizes with TIA1-positive stress granules (lower panel). The distribution of measures from n=3 independent experiments is reported, with each biological replicate color-coded: the mean value from each of the three replicates is represented by black dots, and the mean ± SD of the three replicates is shown as a black line. Statistical significance was calculated by One-way ANOVA test, and the significant differences between the A2 variants or between the A2b variants are shown. *p<0.05, **p<0.01, ****p<0.0001. (D) Representative western blot and relative quantification of SH-SY5Y cells untreated (NT) or transfected with an empty plasmid (mock), or with plasmids coding for HA-tagged hnRNP A2 (A2) and HA-tagged hnRNP A2b (A2b). After 48 h, total protein extracts (input), as well as the nuclear and cytosolic fractions were analysed by western blot with an anti-cleaved PARP (cPARP) and anti-cleaved Caspase 3 (cCasp3) antibodies. A2 and A2b expression levels were monitored using an anti-HA antibody. Lamin B1 and β-actin levels were monitored to assess the fractions’ purity and normalize the fractions. Data are reported as mean value ± SD (n=3 independent experiments). Statistical significance was calculated by One-way ANOVA test, *p<0.05, **p<0.01, ****p<0.0001. (E) Representative western blot and relative quantification of SH-SY5Y cells untreated (NT) or transfected with an empty plasmid (mock), or with plasmids coding for HA-tagged hnRNP A2 (A2), hnRNP A2b (A2b), hnRNP A2_ΔNLS (A2_ΔNLS) and hnRNP A2b_ΔNLS (A2b_ΔNLS). After 48 hrs, cytosolic protein extracts were analyzed in western blot with an anti-cCasp3 antibody. Expression of hnRNP A2/B1 isoforms has been evaluated in total lysates (input) with an anti-HA antibody. β actin antibody was used as a loading control. Data are reported as mean value ± SD (n=3 independent experiments). Statistical significance was calculated by One-way ANOVA test, *p<0.05, **p<0.01, ***p<0.001.

### Functional interaction between FUS and hnRNPA2/B1

Previous data suggests that FUS and hnRNP A2/B1 functionally interact, that FUS-related pathogenic conditions affecting this interaction might contribute to the disease, and that splicing alterations in hnRNP A2/B1 might be involved in this process. To further support these hypotheses, we performed a pilot comparison of datasets from FUS depleted/KO mice and mice depleted of hnRNP A2/B1 ^10,19,28^ (Supplementary Table 2). As illustrated in Figure 7A, a significant overlap exists between splicing events controlled by FUS or hnRNP A2/B1. To lead on these results, we performed a bioinformatic analysis of a set of these common splicing alterations and tested by RT-PCR analysis to which extent these splicing alterations are reproduced in hFUS transgenic animals. Using the Enrichr analysis tool ^29^, we analysed the common 52 splicing events shared by hnRNP A2/B1 downregulation by ASO and FUS downregulation (ASO)/ knock-out (KO). As shown in Figure 7B, Gene Ontology analysis shows that the most significant pathways are related to the control of RNA processing, including RNA splicing, supporting the existence of a cascade of splicing alterations mediated by FUS-hnRNP A2/B1 interaction in the pathogenesis of FUS-mediated ALS. Finally, to validate the targets identified, 11 out of the 22 genes whose alternative splicing is affected in both FUS KO and hnRNP A2/B1 ASO were analysed in extracts from spinal cords of end-stage hFUS mice and age-matched non-transgenic animals. The majority (9/11) are indeed modified in affected animals, with 8 of them reaching statistical significance (Figure 7C). Interestingly, the alternative splicing of hnRNP A2/B1 target genes is affected already at a symptomatic phase of the disease in hFUS transgenic mice, along with the modulation of hnRNP A2/B1 (Supplementary Figure 6). These results support the hypothesis that FUS and hnRNP A2/B1 lie on the same molecular pathway and a coordinated process of alternative splicing changes mediated by FUS and hnRNP A2/B1 is associated to disease progression.

**Figure 7.**
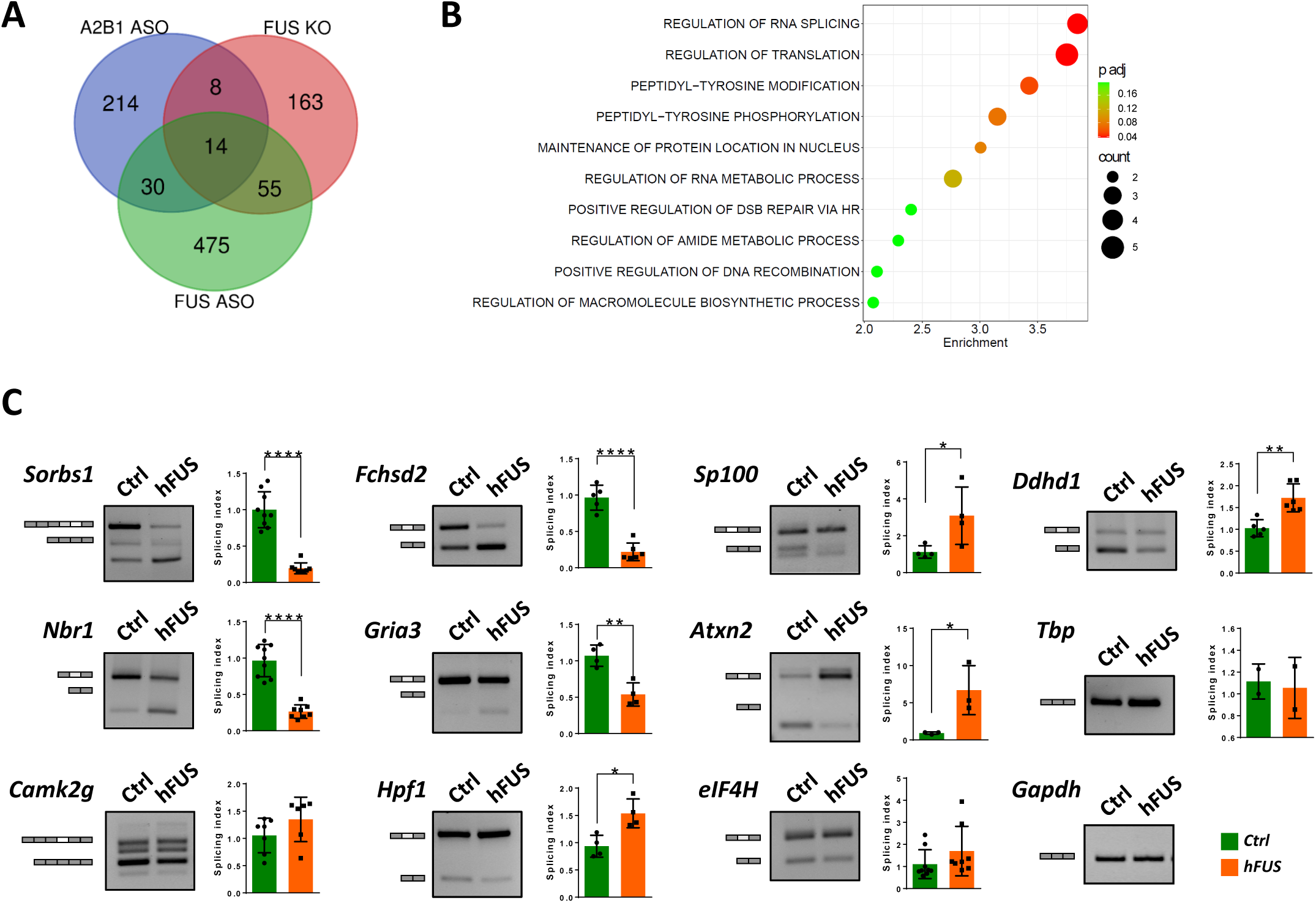
Functional interaction between FUS and hnRNP A2/B1. (A) The Venn diagram illustrates the number of shared alternative splicing changes following FUS depletion (FUS ASO), hnRNP A2/B1 depletion (A2/B1 ASO), or FUS knockout (FUS KO), obtained through a comparative analysis of publicly available datasets ^10,19,28^. (B) Overlapping genes whose alternative splicing is misregulated upon A2/B1 and FUS downregulation (ASO)/ knock-out (KO) have been analysed by the Enrichr analysis tool. The top 10 terms enriched in Gene Ontology biological process are listed according to their decreasing -log10 (p value) (enrichment). The colour code indicates the adjusted p value, and the bubble size reflects the number of genes enriching that annotation (count). (C) The alternative splicing pattern of 11 selected common hnRNP A2/B1 and FUS target genes was assessed through semiquantitative RT-PCR analysis in the spinal cords of end-stage hFUS mice, along with age-matched non-transgenic control (Ctrl) animals. Representations of constitutive exons (dark grey rectangles) and alternatively spliced exons (light grey and white rectangles) analysed are shown. Bands were quantified by densitometric analysis, and a splicing index was calculated as follows. For genes that are expressed in more than one isoform, the ratio between the upper and the lower band was calculated and plotted considering the corresponding ratio in Ctrl mice equal to 1. For genes that are expressed as unique isoform, the splicing index was calculated as the ratio between band intensity and the relative intensity of the housekeeping Gapdh gene. Data are expressed as means ± SD (n≥3mice/group). Statistical significance was calculated by student’s t-test, *p<0.05, **p<0.01, ***p<0.001, ****p<0.0001.

### Splicing switching oligonucleotides promoting exon 9 skipping enhance ALS pathology in hFUS mice

Finally, to assess if the accumulation of hnRNP A2/B1 isoforms lacking exon 9 have a role in the pathogenic process occurring in hFUS mice, we devised splicing switching oligonucleotides (SSOs) to promote exon 9 skipping from hnRNP A2/B1 mRNA. To this aim, a (2’OMe, PS)-modified nucleotide (SSO A) targeting the donor splice sites at the 5’ end defining exon 9 was tested in mouse NSC-34 cells and human HeLa cells. As shown in Supplementary Figure 7A, SSO A induces a significant skipping of exon 9 from hnRNP A2/B1. Under this condition, the accumulation of A2b and B1b protein isoforms is readily detectable by the anti-exon 8/10 antibody (Supplementary Figure 7B), prompting us to assess *in vivo* the effects of an intracerebroventricular (ICV) injection of SSO A on disease phenotypes in hFUS mice. As shown in Supplementary Figure 8, injection of SSO A at post-natal day 1 enhances exon 9 skipping in both spinal cord and cortex, as shown at day 7 and lasting for the overall disease course, with a concurrent increase in hnRNP A2b/B1b protein expression. Significantly, hFUS mice treated with SSO A display an enhanced motoneuron loss (Fig. 8A-B). In the same conditions, SSOA treatment does not significantly affect astrocytosis (Supplementary Figure 9), while it enhances spinal cord microgliosis, as indicated by increased expression levels of Iba1-positive cells, which furthermore display larger area and perimeter compared to hFUS mice treated with vehicle only (Fig 8C-D). Control mice treated with SSOA display neither signs of motor neuron toxicity nor microgliosis (Fig. 8A-C). To deeply investigate the effect of SSOA on microgliosis and infiltrating immune cells, we evaluated the levels of CD68 and P2Y12 proteins, showing that although SSOA treatment does not change the expression of CD68 reactive microglia/macrophages, it significantly decreases the expression of P2Y12 homeostatic microglia (Fig. 8E) in the ventral horn grey matter. Finally, alterations in the alternative splicing of FUS/hnRNP A2/B1 targets recorded in hFUS mice are enhanced by SSOA treatment (Fig. 8F), further indicating that SSOA worsens pathological phenotypes associated to disease in hFUS mice.

**Figure 8.**
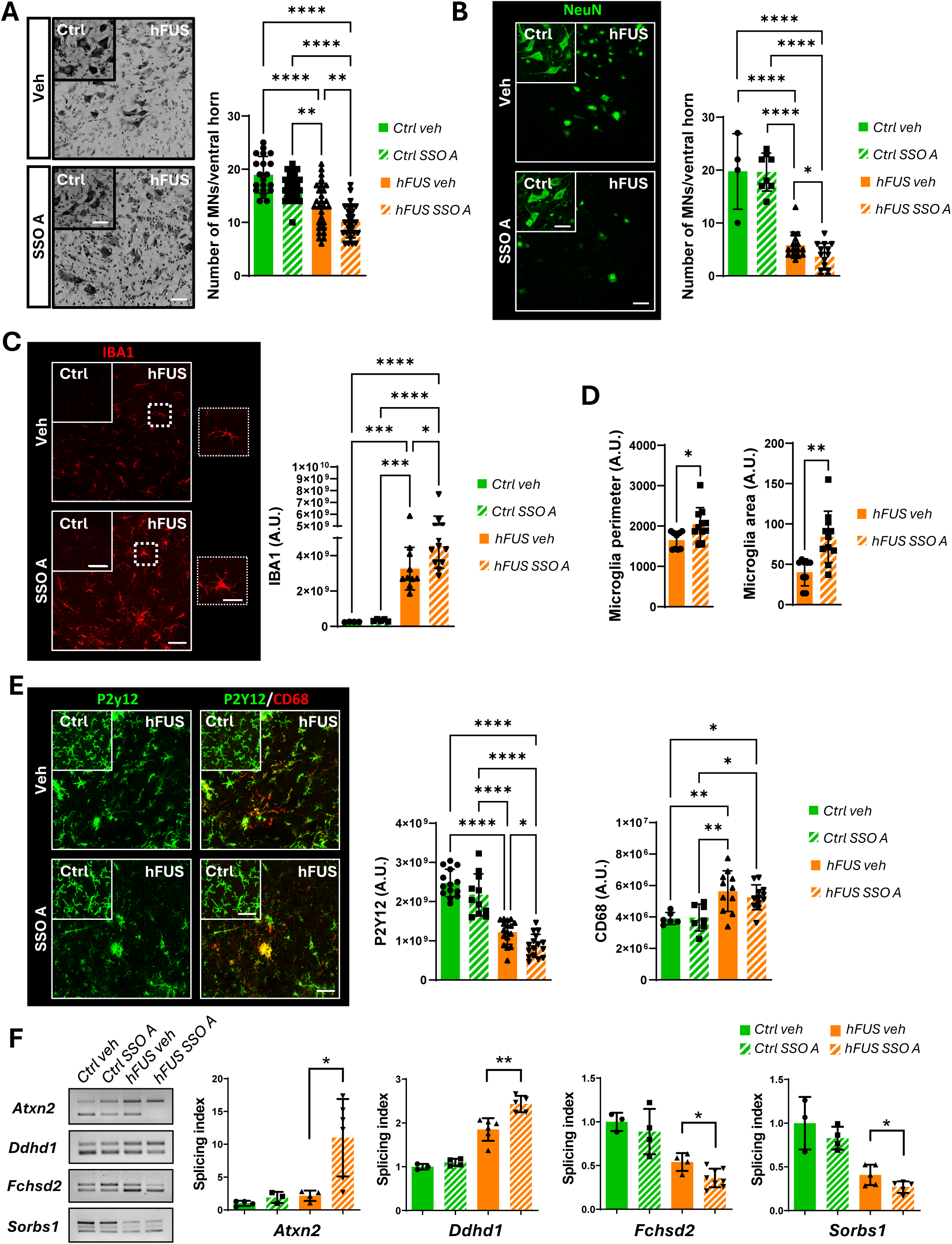
SSO A promotes motor neuron degeneration and enhances neuroinflammation in hFUS mice. (A) Nissl-stained spinal cord sections of non-transgenic (Ctrl) and symptomatic hFUS mice treated with PBS (Veh) or 40 µg SSO A were analyzed 35 days after ICV injection. Scale bar: 50 μm. Quantification of motor neuron (MNs) numbers/ventral horn is provided. Data represent means ± SD. Statistical significance was calculated using ANOVA, **p<0.01, ****p<0.0001. (n = 3/4 animals for group, at least four sections for animal). (B, C, E) Spinal cord sections from non-transgenic (Ctrl) and symptomatic hFUS mice treated with PBS (Veh) or 40 µg SSO A were analyzed 35 days after ICV injection and subjected to immunofluorescence staining with an antibody against NeuN (green) (B), Iba1 (red) (C), P2Y12 (green) and CD68 (red) (E). Scale bar: 50 μm, 4 μm (insert). Data represent means ± SD. Statistical significance was calculated using ANOVA, **p<0.01, ****p<0.0001. (n = 3/4 animals for group, at least four sections for animal). (D) Iba1 positive cells in vehicle- and SSO A treated hFUS mice were analyzed by ImageJ software for different size descriptors (Area and Perimeter,). Data represent means ± SD. Statistical significance was calculated using Student’s t-test, *p<0.05, **p<0.01, (n = 3/4 animals for group, at least four sections for animal). (F) The alternative splicing pattern of Atxn2, Ddhd1, Fchsd2 and Sorbs1 target genes was assessed through semiquantitative RT-PCR analysis in the spinal cords of non-transgenic (Ctrl) and symptomatic hFUS mice treated with PBS (Veh) or 40 µg SSO A. Bands were quantified by densitometric analysis, and a splicing index was calculated as the ratio between band intensity. Data are expressed as means ± SD (at least n=3 mice/group). Statistical significance was calculated using unpaired t-test, *p<0.05, **p<0.01.

## Discussion

The pathogenesis of ALS has been linked to the mislocalization of RBPs from the nucleus, where they normally reside, to the cytoplasm, where they typically form condensates that are supposed to be toxic by a gain of function mechanism ^30,31^. This is clearly established for TDP-43, which is depleted from cell nuclei and accumulates into pathological inclusions in the vast majority of ALS cases ^5,32^, as well as for ALS-linked FUS mutations, that cause incorrect nuclear-cytoplasmic localization and condensation of FUS into cytoplasmic SGs ^33,34^. Importantly, this seems to be also the case for a plethora of other RBPs, that are affected in their expression and distribution in ALS, despite a clear genetic link to the disease has been established only for a few of them ^7,35,36^. Thus, the functional network connecting RBPs is widely affected by ALS conditions, and an extensive process of mislocalization and/or aggregation of RNA-binding proteins (RBPs) can occur along with the progression of the disease and might have significant implications for the overall pathological process.

Our data point to hnRNP A2/B1, an RBP with a role in various steps of RNA metabolism, as a relevant player in this process. Indeed, in mice overexpressing human wild-type FUS (hFUS), which display ALS-related pathological features ^16,37,38^, a major alteration in the mRNA splicing of hnRNP A2/B1 occurs. We observed a significant skipping of exon 9 from the mature RNA, leading to changes in the relative expression of the four known mRNA isoforms encoding hnRNP A2/B1. These splicing alterations arise at the onset of pathological symptoms and persist until the end stage of the disease. Conversely, no variations are observed during the pre-symptomatic phase, supporting the idea that the changes of hnRNP A2/B1 alternative splicing may be associated with the onset/progression of the disease rather than representing a secondary effect associated to the terminal phase. Importantly, our results revealed that the observed changes in alternative splicing correlate with the shift in the expression of hnRNP A2/B1 protein isoforms. Indeed, spinal cord from hFUS mice shows a decreased expression of exon 9-containing isoforms (A2 and B1) with a concomitant increase in isoforms lacking exon 9 (A2b and B1b), in both symptomatic and end-stage phases. Additionally, our results show that under physiological conditions hnRNP A2/B1 isoforms are ubiquitously present across various cell types in the spinal cord, including neuronal and non-neuronal cells. Under pathological conditions, however, a significant reduction in the overall expression of exon 9-containing isoforms occurs, while exon-9 lacking isoforms are readily detectable in degenerating motor neurons, as well as in astrocytes and microglia cells that populate the grey matter of end stage animals, suggesting a role of these isoforms in the pathological process. Most importantly, our study reveals that A2b/B1b isoforms relocate to the cytoplasm of degenerating motor neurons in hFUS mice. Thus, changes in isoform expression of hnRNP A2/B1 in hFUS mice resemble what has been already established for other ALS-associated RBP, that in pathological conditions undergo decreased expression and/or cytoplasmic accumulation ^35,39,40^.

The altered expression of hnRNP A2/B1 may certainly result in a loss of nuclear functions of this protein. Indeed, it is documented that depletion of hnRNP A2/B1 in the mouse spinal cord induces changes in alternative splicing of several target genes ^28^. Interestingly, a significant number of genes that are regulated by hnRNP A2/B1 display alterations in their splicing patterns in the spinal cord of symptomatic and end-stage hFUS mice, suggesting that a loss of splicing regulation by hnRNP A2/B1 has a role in this process. Whether these changes are caused by a decreased nuclear pool of hnRNP A2/B1 or by the concurrent cytoplasmic mis-localization of exon 9-lacking isoforms is still to be fully defined. However, SSOs that increase exon 9 skipping in hFUS mice enhance the observed splicing alterations, suggesting that these changes are tightly linked to A2b/B1b accumulation and to the overall disease progression.

Our analysis revealed that isoforms lacking exon 9 localize within cytoplasmic SGs, and that under conditions known to promote SGs assembly, SG accumulation of A2b/B1b variants is significantly enhanced. Recent data pointed out an important role of hnRNP A2/B1 in regulating SG dynamics ^24^, which is known to be affected in ALS-related conditions ^41–46^, and showed that modifying hnRNP A2/B1 expression affect the dynamics of pathological SGs induced by mutant FUS, thus supporting the hypothesis that A2b/B1b isoforms might impact on this function. Interestingly, we observed that the hnRNP A2 variant lacking a portion of the M9 nuclear localization signal accumulates into SGs, independently from the presence of exon 9, indicating that the accumulation of hnRNP A2/B1 in SGs is a direct consequence of its cytoplasmic delocalization. Moreover, the results obtained from *in vitro* analysis clearly demonstrate that exon 9 plays a crucial role in determining the nuclear-cytoplasmic localization of hnRNP A2/B1 and identify the isoforms lacking exon 9 as the primary cytoplasmic variants of hnRNP A2/B1. This conclusion is supported by previous observations showing that A2b is a predominant cytoplasmic isoform in neural cells, suggesting a potential role in axonal mRNA trafficking regulation ^47^. Exon 9 is situated in proximity to the M9-NLS, and it is possible that the absence of exon 9 may alter the NLS recognition by the nuclear import receptor karyopherin β2, enhancing the cytoplasmic localization of hnRNP A2/B1 isoforms lacking exon 9, which might in turn facilitate their recruitment into pathological inclusions. Importantly, recent evidence highlights that frameshift variants of hnRNP A2/B1, associated with a specific form of muscular dystrophy, modify the C-terminal region of the nuclear localization signal (NLS) in hnRNP A2 ^27^. This alteration promotes the cytoplasmic accumulation of the protein and its recruitment into SGs by disrupting its interaction with karyopherin β2 and in turn causes cell toxicity ^27^, suggesting that uncontrolled cytoplasmic expression of hnRNP A2/B1 might be harmful to cells. A D290V mutant form of hnRNP A2/B1, known to induce motor neuron degeneration in rare cases of familial ALS, also undergoes re-localization from the nucleus to the cytoplasm, where it accumulates in an aggregated and insoluble form ^28^. In addition, muscle biopsies from patients associated with multisystem proteinopathy (an autosomal dominant disease with features similar to ALS) carrying the D290V mutation showed that hnRNP A2/B1 was cleared from the nuclei of fibers and accumulated in cytoplasmic inclusions ^23^. Although mutations in hnRNP A2/B1 provide a minor contribution to genetic ALS cases ^23,28^, our findings indicate that conditions promoting cytoplasmic delocalization of hnRNP A2/B1 might have significant consequences for cellular viability, and suggest that the cytoplasmic accumulation of A2b/B1b isoforms that we recorded in hFUS mice might play an important role specifically in motor neuron degeneration. This adds to previous findings showing that cytoplasmic accumulation of pathogenic FUS causes an imbalance in the homeostasis RBPs, including hnRNP A2/B1, that exacerbates neurodegeneration ^48^.

Results from cultured neuronal cells support the conclusion that an increased cytoplasmic expression of hnRNP A2/B1 might be harmful to cells. Indeed, cytoplasmic hnRNP A2b isoform promotes a significant increase in activated caspase-3 expression compared to controls and to cells expressing A2. Interestingly, the A2_ΔNLS isoform, which, despite containing exon 9, relocates to the cytosol upon removal of the M9_NLS, induces caspase-3 activation, demonstrating that uncontrolled cytoplasmic delocalization of A2 is sufficient to promote cellular toxicity. This further suggests that the exclusion of exon 9, and the subsequent cytoplasmic accumulation of the A2b isoform, may play a role in the pathogenic mechanism of FUS-related ALS. The experiments performed by *in-vivo* injection of (SSOs) strengthen this conclusion. Indeed, enhanced exon 9 skipping induced by SSO treatment increases motor neuron degeneration and neuroinflammation that characterize the disease course in hFUS mice, demonstrating that the accumulation of A2b/B1b isoforms contributes to FUS-associated toxicity in mice.

Overall, our findings support the notion that ALS conditions broadly impact the functional network of RNA-binding proteins and identify hnRNP A2/B1 mislocalization as a possible player in the pathological process characterizing FUS-ALS.

## Materials and Methods

### Animal model

Adult Tg (Prnp-FUS) WT3Cshw/J mice expressing hemagglutinin-tagged human wild-type FUS (hFUS) were obtained from Jackson Laboratories. Mice were maintained in hemizygosity on the same C57BL/6 genetic background. Hemizygous FUS mice were backcrossed to obtain homozygous mice, used as experimental subjects. Animals were housed in our indoor animal facility at constant temperature (22 ± 1 °C) and relative humidity (50%) with 12-h light cycle (light 7 am–7 pm). Food and water were freely available. When animals showed symptoms of paralysis, wet food was given daily into the cages for easy access to nutrition and hydration. All animal procedures were performed according to the European Guidelines for the use of animals in research (2010/63/EU) and the requirements of Italian laws (D.L. 26/2014). The ethical procedure was approved by the Italian Ministry of Health (protocol number 383/2022 PR/G). Every attempt was made to reduce animal suffering and minimize the required number of animals to ensure the generation of reliable results. ALS-like disease stages were determined using a neurological score assessment ^49^. Animals were assigned at the presymptomatic phase at 20 days old, with no clinical signs of disease. The symptomatic phase was assigned to homozygous transgenic mice (hFUS) aged 32-34 days, when mice showed a reduction in locomotor activity compared to non-transgenic control (Ctrl) and hemizygous mice, displaying signs of start hind limb paralysis. End-stage animals were sacrificed at 38-40 days old, when the atrophy of both hind limbs was detected. Mice were genotyped by PCR analysis of tissue extracts from tail tips, as previously described ^38^.

### Plasmids and Splicing Switching Oligonucleotides (SSOs)

Plasmids containing degenerated protein-coding sequences of wild type and mutated D290V hnRNP A2 isoforms were purchased by Addgene. hnRNP B1 sequence were amplified by PCR from hnRNP A2 plasmid. All other variants, including hnRNP B1b, A2b and A2b D290V isoforms, were produced by PCR-driven overlap extension. Δ323-341 deletion variants (hnRNP A2_ΔNLS and hnRNP A2b ΔNLS) were mutagenized by PCR from wild type hnRNP A2 and hnRNP A2b plasmids. An extra canonical SV-40 nuclear localization signal has been introduced in wild type hnRNP A2 and hnRNP A2b plasmids by PCR to produce hnRNP A2+NLS and hnRNP A2b+NLS variants. All hnRNP A2/B1 variants were cloned into pcDNA3.1 plasmid vector (Invitrogen) and fused with an HA epitope at the N-terminal end.

Fully modified 2’-O-methyl splicing switching oligonucleotides (SSO) with a phosphorothioate backbone have been synthesized and HPLC purified by Eurofins Genomics for subsequent test in vitro. HPLC purified and dialyzed, with low endotoxin level (guaranteed < 0.5 EU/mg), SSO was synthesized by Microsynth AG for studies in vivo. The sequences of SSOs used are: SSO A 5’-UUUAUUACCUCCUCCA-3’ and scramble SSO 5’-ACCUUCUUACUUCAUC-3’.

### Cell cultures, treatment and transfection

Human HeLa and SH-SY5Y, and mouse NSC34 cells were cultured in Dulbecco’s modified Eagle’s medium (DMEM) with Glutamax (Corning), supplemented with 10% fetal bovine serum (FBS, Euroclone), and 1% penicillin/streptomycin (Sigma-Aldrich) at 37 °C in a 5% CO2 atmosphere. To induce the integrated stress response, cells were treated with sodium arsenite (NaAs, Sigma Aldrich) at a concentration of 0.5 mM. For transient transfection experiments, cells at 80% confluence were transfected with appropriate plasmids or antisense oligonucleotides using Lipofectamine 2000 (Invitrogen) according to the manufacturer’s instruction. Cells were collected after 24 and/or 48 hours for subsequent analysis.

### Cortical neuron primary cultures

Primary cortical neuron cultures were isolated from the cerebral cortex of postnatal C57BL/6 mice (P0-P1) following established protocols ^50,51^. Mice were decapitated, and after meningeal removal, cortices were minced, digested with 0.25% trypsin (Gibco) and 0.2 mg/ml DNase (Sigma-Aldrich) in DMEM, and incubated at 37°C for 20 minutes. After dissociation with a fire-polished Pasteur pipette and passage through 70 μm filters, primary cortical neurons were suspended and plated in 12-well plates previously coated with poly-L-lysine (PLL) (1 mg/mL) or on PLL-coated coverslips and maintained in Neurobasal® medium (Gibco Life Technologies) supplemented with B-27® (Life Technologies) at densities ranging from 4 x 10^4^ cells/cm^2^ to 6 x 10^4^ cells/cm^2^.

### Protein extraction from tissue and cells

Lumbar spinal cord from n=4 animals per group were dissected and homogenized with a homogenizer (MICCRA D-1) in lysis buffer containing 20 mM Hepes pH 7.4, 100 mM NaCl, 1% Triton, 10 mM EDTA and a protease inhibitor cocktail (Sigma-Aldrich). After an incubation of 45 minutes in ice, the lysates were centrifugated for 20 minutes at 16000 × g at 4°C. The supernatant was quantified using the Bradford assay and resuspended in Laemmli buffer (Biorad). SH-SY5Y and Hela cells were lysed in RIPA buffer (50 mM Tris-HCl pH 7.4, 1% Triton, 0.25% sodium deoxycholate, 0.1% SDS, 150 mM NaCl, 1 mM EGTA, 5 mM MgCl2) containing a protease inhibitor cocktail, incubated for 30 minutes on ice and centrifugated for 10 minutes at 16000 × g at 4°C. Supernatants were quantified using the Bradford protein assay (Bio-Rad) and resuspended in Laemmli buffer. For the preparation of insoluble protein extracts, the pellets were resuspended in Laemmli Buffer.

### Nuclei-cytoplasm fractionation

For subcellular fractionation, HeLa cells were centrifuged at 600 × g for 5 minutes at 4 °C and washed with cold PBS. Cell pellet was resuspended by gentle pipetting with cold hypotonic lysis buffer (HLB, Tris 10 mM pH 7.5; NaCl 10 mM; MgCl2 3 mM; NP-40 0.1%; glycerol 10%), with 1 mM sodium orthovanadate, 1 mM sodium fluoride and a cocktail of protease inhibitors (Sigma-Aldrich). After an incubation on ice for 10 minutes, Laemmli buffer was added to a portion of lysates to obtain the total fraction. The remnant cell suspension was centrifuged at 1000 x g for 3 minutes at 4 °C. Supernatant containing the cytoplasmic fraction was clarified at 5000 × g for 5 minutes at 4 °C, quantified by Bradford assay and then resuspended in Laemmli Buffer. Pellet containing the nuclear fraction was washed by carefully pipetting with cold PBS, centrifuged at 300 × g for 2 minutes at 4°C, and then resuspended in Laemmli Buffer.

### SDS-PAGE and western blotting

Protein lysates were separated by SDS-PAGE and transferred to a nitrocellulose membrane. The membranes were blocked at room temperature for 1 hour in Tris-buffered saline solution with 0.1% Tween-20 (TBS-T) containing 5% non-fat dry milk, and then incubated with primary antibodies, diluted in TBS-T containing 2% non-fat dry milk, at 4°C overnight or for 2 hours at room temperature. After washing, HRP-conjugated secondary antibodies (Jackson ImmunoResearch) were applied at room temperature for 1 hour. Chemiluminescent detection was performed using ECL solution (Roche). Following densitometry-based quantification and analysis using ImageJ software (National Institute of Health, NIH), the relative density of each identified protein was calculated.

### Immunofluorescence analysis and confocal microscopy

Spinal cords from n=4 animals per group were fixed using a 4% paraformaldehyde solution (PFA) in 0.1M PBS for 12 hours and tissues were cryoprotected in 30% sucrose in PBS solution at 4°C. Subsequently, spinal cords were cut into 30-μm-thick slices with a freezing cryostat (Leica Biosystems). After blocking for 1 hour in 10% normal donkey serum (NDS) in PBS containing 0.3% Triton X-100, spinal cord slices were incubated for 3 days at 4 °C with primary antibodies diluted in 2% NDS in PBS, 0.3% Triton X-100, and then for 3 h at room temperature with appropriate fluorescent secondary antibody, diluted in the same solution. Nuclei were stained with 1 μg/ml DAPI (Sigma-Aldrich) for 10 min. The slides were coverslipped with Fluromount Aqueous Mounting Medium (Sigma-Aldrich). Primary cortical neurons, HeLa and SH-SY5Y cells were fixed using 4% PFA in PBS for 10 minutes, permeabilized with a 0.1% Triton X-100 solution in PBS for 5 minutes and blocked with 2% FBS diluted in PBS for 30 minutes at room temperature. Cells were then incubated with primary antibodies diluted in 2% FBS in PBS for 1 hour at 37° and, subsequently, with appropriate fluorescent-conjugated secondary antibodies in PBS, 2% FBS for 1 hour at room temperature. Nuclei were stained with 1 μg/ml DAPI (Sigma-Aldrich) for 5 min. Immunofluorescence images were analysed using a LEICA TCS SP5 confocal microscope. Images were captured under constant exposure time, gain, and offset settings. Digital image brightness and contrast were adjusted using the LAS AF software (Leica). Background subtraction was performed after defining a region of interest, and the average pixel intensity was calculated. All image quantifications were done using ImageJ software (NIH). For the quantification of cells displaying cytoplasmic and stress granule localization of HA-hnRNP A2/B1 isoforms, at least 50-100 HA-positive cells per condition from randomly selected fields in n=3 independent experiments were visually scored using a Zeiss Axioplan fluorescence microscope.

### Nissl staining

The total number of motor neurons in the L3–L5 segments of the lumbar spinal cord was quantified by analyzing serial sections from each mouse. Sections were cut at a thickness of 30 μm. To visualize the Nissl substance within neurons, the sections were stained with 0.02 % cresyl violet solution. Following staining, the sections underwent a graded dehydration process using ethanol (50% to 100%), were cleared with xylene, and mounted using Eukitt (Sigma-Aldrich, Saint Louis, MO, USA). Images of the sections were captured using a Zeiss Axioskop 2 microscope at 20x magnification. Both the right and left ventral horns were examined to count neurons, characterized by cell bodies exceeding 200 μm², and the average count from the sections was calculated for each mouse.

### Antibodies

Immunofluorescences (IF) and immunoblots (WB) were performed with the following primary antibodies: rabbit anti-Exon 2 (1:1000-WB, 1:500-IF), rabbit anti-Exon 9 (1:1000-WB, 1:500-IF) and rabbit anti-Exon 8/10 (1:500-WB, 1:100-IF) were kindly provided by Prof. Smith and Prof. Rothnagel from the University of Queensland (Australia); rabbit anti-Exon 8/10 (1:5000-WB, 1:5000-IF, custom antibody produced by Bio-Fab Research), rabbit anti-pan hnRNP A2/B1 (1:5000-WB, GeneTex), mouse anti-SMI32 (1:1000-IF, BioLegend), mouse anti-GFAP (1:1000-IF, Cell Signaling), rat anti-CD11b (1:200, Bio-Rad), rabbit anti-β3-Tubulin (1:1000-IF, Cell Signaling), anti-HA mouse (1:2000-WB, 1:500-IF, Sigma Aldrich), goat anti-TIA1 (1:200-IF, Santa Cruz Biotechnology), rabbit anti-cleaved Caspase 3 (1:1000-WB, Cell Signaling Technology), rabbit anti-Lamin B1 (1:3000-WB, Abcam), rabbit anti-cleaved Parp (1:1000-WB, Cell Signaling Technology), mouse anti-GAPDH (1:10000-WB, Calbiochem), mouse anti-β-actin (1:5000-WB, Sigma-Aldrich), rabbit anti-Iba1 (1:500-IF, Wako), rat anti-CD68 (1:500-IF, Biorad), rabbit anti-P2Y12 (1:200-IF, Anaspec), rabbit anti-NeuN (1:500-IF, Cell Signaling).

Secondaries antibodies for WB were anti-rabbit (1:2500) and anti-mouse (1:5000) IgG peroxidase-conjugated from Bio-Rad Laboratories (Hercules, CA, USA). Secondary fluorescent antibodies for IF were Alexa-Flour 488-Donkey anti-rabbit (1:200), Cy3-Donkey anti-mouse (1:200), Cy3-Donkey anti-rat (1:200), Alexa-Flour 488-Donkey anti-goat (1:200), from Jackson ImmunoResearch Laboratories (West Grove, PA, USA).

### Semiquantitative RT-PCR

RNAs were extracted using Trizol reagent (Invitrogen), treated with DNase I (Promega), and reverse transcribed using the ImProm-II™ Reverse Transcription System (Promega), following the manufacturer’s instructions. Semiquantitative RT-PCR reactions were performed using Biomix Red (Bioline) using 50 ng cDNA and 0,5 µM of specific primers, listed in Supplementary Table 1. PCR products were run in 2% agarose gels and visualized by SYBR Safe DNA Gel Stain (Invitrogen) staining. Images were acquired on ChemiDocTM Imaging System (Bio-Rad), bands were quantified using the ImageJ software (NIH) and the splicing indices were calculated as the ratio between the upper and the lower bands. The expression levels of the isoform lacking exon 9 were calculated as percentages relative to the total expression of the isoform containing exon 9 and the one lacking exon 9.

### Granule Protein Enrichment Server (Grapes)

The information pertaining to the propensity of hnRNP A2 and hnRNP A2b to localize into cellular condensates were obtained by the prediction tool Granule Protein Enrichment Server (Grapes) ^26^, using the MaGSeq (MaGS Sequence-based tool) predictive model. MaGSeq is a general linearized model (GLM) based only on protein sequence features and provides a Z-score, representing the propensity of proteins to localize into biological condensates, as well as the feature scores used to generate the predictions. A MaGSeq value greater than or equal to 0.90 for human suggests that the protein is highly likely part of phase-separated organelles within the cell.

### Intracerebroventricular injection of SSO

Sterile PBS or SSO A (20 μg or 40 μg) diluted in 0.01% Fast Green (Sigma-Aldrich) was injected intracerebroventricularly (ICV) in newborn pups (P0-P1) using a glass syringe (Hamilton, model 75 RN 5 μL Syringe). P0-P1 mice were anesthetized on ice, and for the single ICV injection, the needle was inserted at the midpoint of a line defined between the right eye and the lambda intersection of the skull. The needle was carefully advanced into the lateral ventricle to a depth of approximately 3 mm. Following the injection, the pups were allowed to recover on a heating pad under a heat lamp before being returned to their mother.

### Statistics

Data are reported as mean ± SD. For comparison between two groups, statistical significance was assessed using a two-tailed Student’s t-test. For multiple comparisons, One-way analysis of variance (ANOVA) or Two-way ANOVA, followed by Tukey’s test, were employed. All statistical analyses were conducted using GraphPad Prism 9.0 software (GraphPad Software, San Diego, CA, USA), with significance set at p<0.05. Animals were randomly used for experiments.

## Acknowledgements

This work was supported by Fondazione Arisla ETS (Project Spliceals to M.C., N.D.A., G.C, and Project SwitchALS to M.C. and N.D.A.), CNR (project Nutrage, IFT DBA.AD005.225 to M.C.), and European Union - Next Generation EU and founded by the Ministry of University and Research (MUR), National Recovery and Resilience Plan (PNRR), project MNESYS (PE0000006) – A Multiscale Integrated Approach to the Study of the Nervous System in Health and Disease (DN. 1553 October 11, 2022). S.R. and A.S. are supported by European Union - Next Generation EU, within the PNRR project “Rome Technopole - Innovation Ecosystem”. Dr Valeria Gerbino (Fondazione Santa Lucia, IRCCS, Rome, Italy) is gratefully acknowledged for providing help with ICV in vivo injection of SSOs. We thank Dr. Joseph Rothnagel (School of Chemistry and Molecular Biosciences, University of Queensland, Australia) for providing hnRNP A2/B1 isoform specific antibodies, and Dr Ivan Arisi, (EBRI Foundation, Rome, Italy) for the initial dataset analysis.

## Ethics declarations

### Ethics approval

All animal procedures were performed according to the European Guidelines for the use of animals in research (2010/63/EU) and the requirements of Italian laws (D.L. 26/2014). The ethical procedure was approved by the Italian Ministry of Health (protocol number 383/2022 PR/G).

### Availability of Data and Materials

Data available on request from the authors

### Contributions

S.R. conceived the idea, designed, performed, and interpreted most of the molecular and cell biology experiments, wrote and edited the paper. M.M. designed, performed, and interpreted most of the mouse biology experiments, and wrote and edited the paper. I.D.V. helped with mouse experiments and ICV injection of SSO. S.B. helped with cell biology experiments and with the design and in vitro testing of SSO. V.D. helped with cell biology experiments. M.A. helped with the molecular analysis of SSO effects. E.D.A. aided with the maintenance of cell cultures and sample generation. M.D.S. aided with sample preparation and analysis. A.S. helped with confocal microscopy analysis. G.C. generated funding and edited the paper. S.A. conceived the idea, designed, supervised and interpreted the experiments, wrote and edited the paper. N.D.A. generated funding, conceived the idea, designed, supervised and interpreted the experiments, wrote and edited the paper. M.C. generated funding, conceived the idea, designed, supervised and interpreted the experiments, wrote and edited the paper.

### Corresponding authors

Correspondence to Savina Apolloni, Nadia D’Ambrosi or Mauro Cozzolino

## References

1 Xue YC, Ng CS, Xiang P, Liu H, Zhang K, Mohamud Y et al. Dysregulation of RNA-Binding Proteins in Amyotrophic Lateral Sclerosis. Front Mol Neurosci 2020; 13: 78.

2 Kim G, Gautier O, Tassoni-Tsuchida E, Ma XR, Gitler AD. ALS Genetics: Gains, Losses, and Implications for Future Therapies. Neuron 2020; 108: 822–842.

3 Nussbacher JK, Tabet R, Yeo GW, Lagier-Tourenne C. Disruption of RNA Metabolism in Neurological Diseases and Emerging Therapeutic Interventions. Neuron 2019; 102: 294–320.

4 Taylor JP, Brown RH, Cleveland DW. Decoding ALS: from genes to mechanism. Nature 2016; 539: 197– 206.

5 Suk TR, Rousseaux MWC. The role of TDP-43 mislocalization in amyotrophic lateral sclerosis. Mol Neurodegener 2020; 15: 45.

6 Mehta PR, Brown A-L, Ward ME, Fratta P. The era of cryptic exons: implications for ALS-FTD. Mol Neurodegeneration 2023; 18: 16.

7 Wolozin B, Ivanov P. Stress granules and neurodegeneration. Nat Rev Neurosci 2019; 20: 649–666.

8 Baradaran-Heravi Y, Van Broeckhoven C, Van Der Zee J. Stress granule mediated protein aggregation and underlying gene defects in the FTD-ALS spectrum. Neurobiology of Disease 2020; 134: 104639.

9 Li YR, King OD, Shorter J, Gitler AD. Stress granules as crucibles of ALS pathogenesis. Journal of Cell Biology 2013; 201: 361–372.

10 Scekic-Zahirovic J, Sendscheid O, El Oussini H, Jambeau M, Sun Y, Mersmann S et al. Toxic gain of function from mutant FUS protein is crucial to trigger cell autonomous motor neuron loss. EMBO J 2016; 35: 1077– 1097.

11 Ishigaki S, Sobue G. Importance of Functional Loss of FUS in FTLD/ALS. Front Mol Biosci 2018; 5: 44.

12 Shang Y, Huang EJ. Mechanisms of FUS mutations in familial amyotrophic lateral sclerosis. Brain Research 2016; 1647: 65–78.

13 Ling S-C, Polymenidou M, Cleveland DW. Converging Mechanisms in ALS and FTD: Disrupted RNA and Protein Homeostasis. Neuron 2013; 79: 416–438.

14 Mackenzie IRA, Neumann M. Fused in Sarcoma Neuropathology in Neurodegenerative Disease. Cold Spring Harb Perspect Med 2017; 7: a024299.

15 Sabatelli M, Moncada A, Conte A, Lattante S, Marangi G, Luigetti M et al. Mutations in the 3’ untranslated region of FUS causing FUS overexpression are associated with amyotrophic lateral sclerosis. Hum Mol Genet 2013; 22: 4748–4755.

16 Ling S-C, Dastidar SG, Tokunaga S, Ho WY, Lim K, Ilieva H et al. Overriding FUS autoregulation in mice triggers gain-of-toxic dysfunctions in RNA metabolism and autophagy-lysosome axis. Elife 2019; 8: e40811.

17 Mitchell JC, McGoldrick P, Vance C, Hortobagyi T, Sreedharan J, Rogelj B et al. Overexpression of human wild-type FUS causes progressive motor neuron degeneration in an age- and dose-dependent fashion. Acta Neuropathol 2013; 125: 273–288.

18 Dini Modigliani S, Morlando M, Errichelli L, Sabatelli M, Bozzoni I. An ALS-associated mutation in the FUS 3′-UTR disrupts a microRNA–FUS regulatory circuitry. Nat Commun 2014; 5: 4335.

19 Lagier-Tourenne C, Polymenidou M, Hutt KR, Vu AQ, Baughn M, Huelga SC et al. Divergent roles of ALS-linked proteins FUS/TLS and TDP-43 intersect in processing long pre-mRNAs. Nat Neurosci 2012; 15: 1488–1497.

20 Mirra A, Rossi S, Scaricamazza S, Di Salvio M, Salvatori I, Valle C et al. Functional interaction between FUS and SMN underlies SMA-like splicing changes in wild-type hFUS mice. Sci Rep 2017; 7: 2033.

21 Humphrey J, Birsa N, Milioto C, McLaughlin M, Ule AM, Robaldo D et al. FUS ALS-causative mutations impair FUS autoregulation and splicing factor networks through intron retention. Nucleic Acids Res 2020; 48: 6889–6905.

22 Rezvykh AP, Ustyugov AA, Chaprov KD, Teterina EV, Nebogatikov VO, Spasskaya DS et al. Cytoplasmic aggregation of mutant FUS causes multistep RNA splicing perturbations in the course of motor neuron pathology. Nucleic Acids Research 2023; 51: 5810–5830.

23 Kim HJ, Kim NC, Wang Y-D, Scarborough EA, Moore J, Diaz Z et al. Mutations in prion-like domains in hnRNPA2B1 and hnRNPA1 cause multisystem proteinopathy and ALS. Nature 2013; 495: 467–473.

24 Wang X, Fan X, Zhang J, Wang F, Chen J, Wen Y et al. hnRNPA2B1 represses the disassembly of arsenite-induced stress granules and is essential for male fertility. Cell Reports 2024; 43: 113769.

25 Wiedner HJ, Giudice J. It’s not just a phase: function and characteristics of RNA-binding proteins in phase separation. Nat Struct Mol Biol 2021; 28: 465–473.

26 Kuechler ER, Jacobson M, Mayor T, Gsponer J. GraPES: The Granule Protein Enrichment Server for prediction of biological condensate constituents. Nucleic Acids Research 2022; 50: W384–W391.

27 Kim HJ, Mohassel P, Donkervoort S, Guo L, O’Donovan K, Coughlin M et al. Heterozygous frameshift variants in HNRNPA2B1 cause early-onset oculopharyngeal muscular dystrophy. Nat Commun 2022; 13: 2306.

28 Martinez FJ, Pratt GA, Van Nostrand EL, Batra R, Huelga SC, Kapeli K et al. Protein-RNA Networks Regulated by Normal and ALS-Associated Mutant HNRNPA2B1 in the Nervous System. Neuron 2016; 92: 780–795.

29 Kuleshov MV, Jones MR, Rouillard AD, Fernandez NF, Duan Q, Wang Z et al. Enrichr: a comprehensive gene set enrichment analysis web server 2016 update. Nucleic Acids Res 2016; 44: W90–97.

30 Bampton A, Gittings LM, Fratta P, Lashley T, Gatt A. The role of hnRNPs in frontotemporal dementia and amyotrophic lateral sclerosis. Acta Neuropathol 2020; 140: 599–623.

31 Purice MD, Taylor JP. Linking hnRNP Function to ALS and FTD Pathology. Front Neurosci 2018; 12: 326.

32 de Boer EMJ, Orie VK, Williams T, Baker MR, De Oliveira HM, Polvikoski T et al. TDP-43 proteinopathies: a new wave of neurodegenerative diseases. J Neurol Neurosurg Psychiatry 2020; 92: 86–95.

33 Bosco DA, Lemay N, Ko HK, Zhou H, Burke C, Kwiatkowski TJ et al. Mutant FUS proteins that cause amyotrophic lateral sclerosis incorporate into stress granules. Hum Mol Genet 2010; 19: 4160–4175.

34 Scekic-Zahirovic J, Sanjuan-Ruiz I, Kan V, Megat S, De Rossi P, Dieterlé S et al. Cytoplasmic FUS triggers early behavioral alterations linked to cortical neuronal hyperactivity and inhibitory synaptic defects. Nat Commun 2021; 12: 3028.

35 Kapeli K, Martinez FJ, Yeo GW. Genetic mutations in RNA-binding proteins and their roles in ALS. Hum Genet 2017; 136: 1193–1214.

36 Akçimen F, Lopez ER, Landers JE, Nath A, Chiò A, Chia R et al. Amyotrophic lateral sclerosis: translating genetic discoveries into therapies. Nat Rev Genet 2023; 24: 642–658.

37 Mitchell JC, McGoldrick P, Vance C, Hortobagyi T, Sreedharan J, Rogelj B et al. Overexpression of human wild-type FUS causes progressive motor neuron degeneration in an age- and dose-dependent fashion. Acta Neuropathol 2013; 125: 273–288.

38 Milani M, Della Valle I, Rossi S, Fabbrizio P, Margotta C, Nardo G et al. Neuroprotective effects of niclosamide on disease progression via inflammatory pathways modulation in SOD1-G93A and FUS-associated amyotrophic lateral sclerosis models. Neurotherapeutics 2024; 21: e00346.

39 Kim W, Kim D-Y, Lee K-H. RNA-Binding Proteins and the Complex Pathophysiology of ALS. Int J Mol Sci 2021; 22: 2598.

40 Zhao M, Kim JR, van Bruggen R, Park J. RNA-Binding Proteins in Amyotrophic Lateral Sclerosis. Mol Cells 2018; 41: 818–829.

41 Hu L, Mao S, Lin L, Bai G, Liu B, Mao J. Stress granules in the spinal muscular atrophy and amyotrophic lateral sclerosis: The correlation and promising therapy. Neurobiology of Disease 2022; 170: 105749.

42 Alirzayeva H, Loureiro R, Koyuncu S, Hommen F, Nabawi Y, Zhang WH et al. ALS-FUS mutations cause abnormal PARylation and histone H1.2 interaction, leading to pathological changes. Cell Reports 2024; 43: 114626.

43 Patel A, Lee HO, Jawerth L, Maharana S, Jahnel M, Hein MY et al. A Liquid-to-Solid Phase Transition of the ALS Protein FUS Accelerated by Disease Mutation. Cell 2015; 162: 1066–1077.

44 Murakami T, Qamar S, Lin JQ, Schierle GSK, Rees E, Miyashita A et al. ALS/FTD Mutation-Induced Phase Transition of FUS Liquid Droplets and Reversible Hydrogels into Irreversible Hydrogels Impairs RNP Granule Function. Neuron 2015; 88: 678–690.

45 Ueda T, Takeuchi T, Fujikake N, Suzuki M, Minakawa EN, Ueyama M et al. Dysregulation of stress granule dynamics by DCTN1 deficiency exacerbates TDP-43 pathology in Drosophila models of ALS/FTD. acta neuropathol commun 2024; 12: 20.

46 Boeynaems S, Bogaert E, Kovacs D, Konijnenberg A, Timmerman E, Volkov A et al. Phase Separation of C9orf72 Dipeptide Repeats Perturbs Stress Granule Dynamics. Molecular Cell 2017; 65: 1044–1055.e5.

47 Han SP, Friend LR, Carson JH, Korza G, Barbarese E, Maggipinto M et al. Differential subcellular distributions and trafficking functions of hnRNP A2/B1 spliceoforms. Traffic 2010; 11: 886–898.

48 Marrone L, Drexler HCA, Wang J, Tripathi P, Distler T, Heisterkamp P et al. FUS pathology in ALS is linked to alterations in multiple ALS-associated proteins and rescued by drugs stimulating autophagy. Acta Neuropathol 2019; 138: 67–84.

49 Hatzipetros T, Kidd JD, Moreno AJ, Thompson K, Gill A, Vieira FG. A Quick Phenotypic Neurological Scoring System for Evaluating Disease Progression in the SOD1-G93A Mouse Model of ALS. J Vis Exp 2015; : 53257.

50 Muramatsu R, Yamashita T. Primary Culture of Cortical Neurons. BIO-PROTOCOL 2013; 3. doi:10.21769/BioProtoc.496.

51 Beaudoin GMJ, Lee S-H, Singh D, Yuan Y, Ng Y-G, Reichardt LF et al. Culturing pyramidal neurons from the early postnatal mouse hippocampus and cortex. Nat Protoc 2012; 7: 1741–1754.

