## Supplementary Material for "Cytoplasmic accumulation of a splice variant of hnRNPA2/B1 contributes to FUS-associated toxicity in a mouse model of ALS"

### Supplementary FIGURE 1

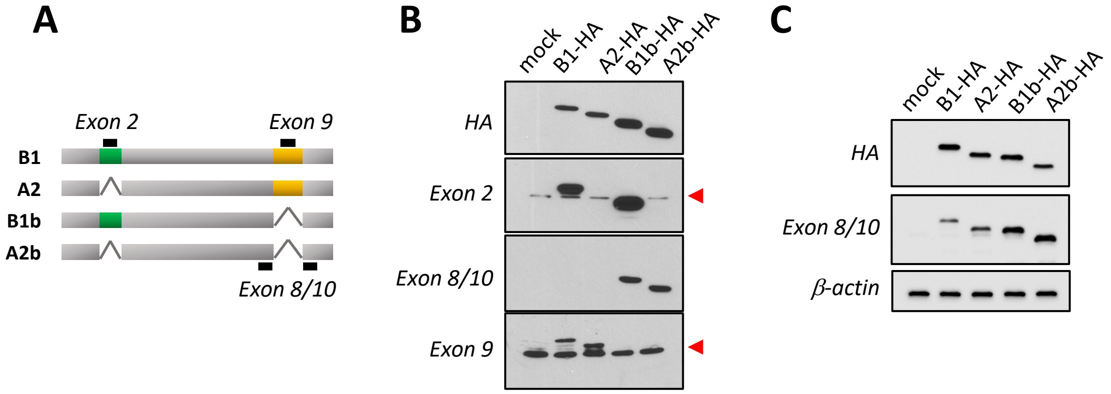

Supplementary FIGURE 2

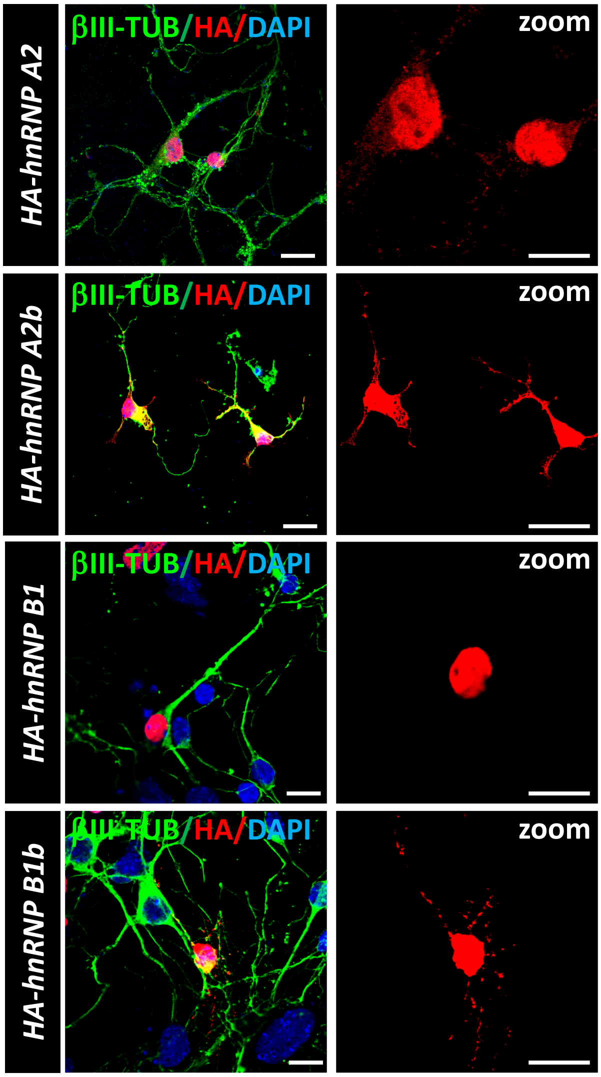

Supplementary FIGURE 3

A

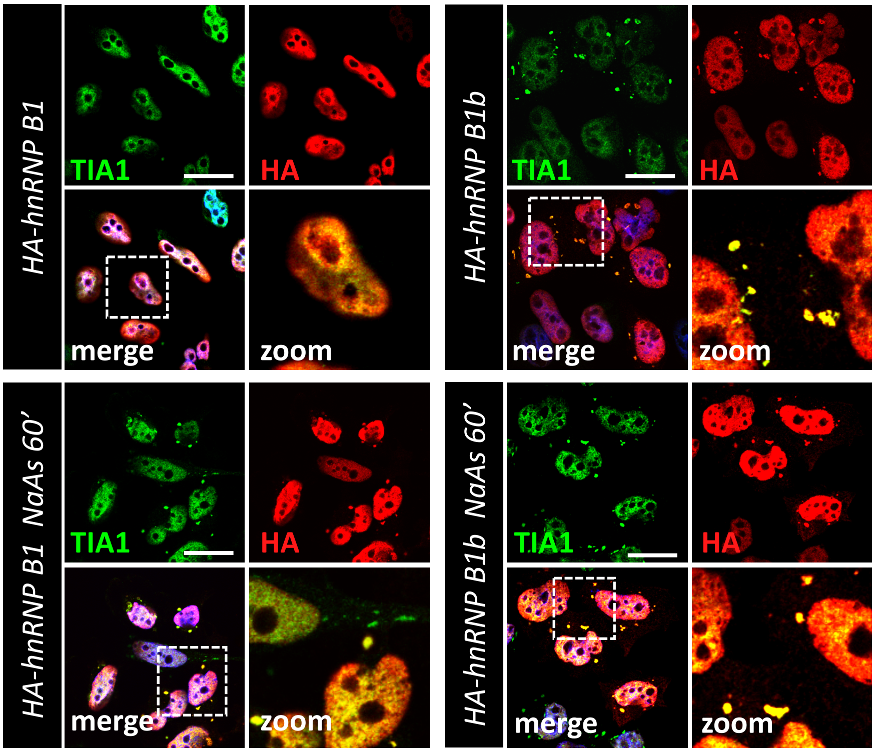


B

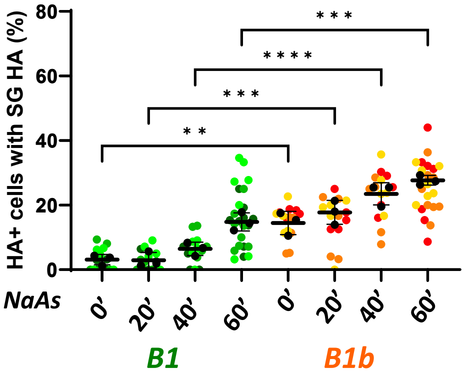

Supplementary FIGURE 4

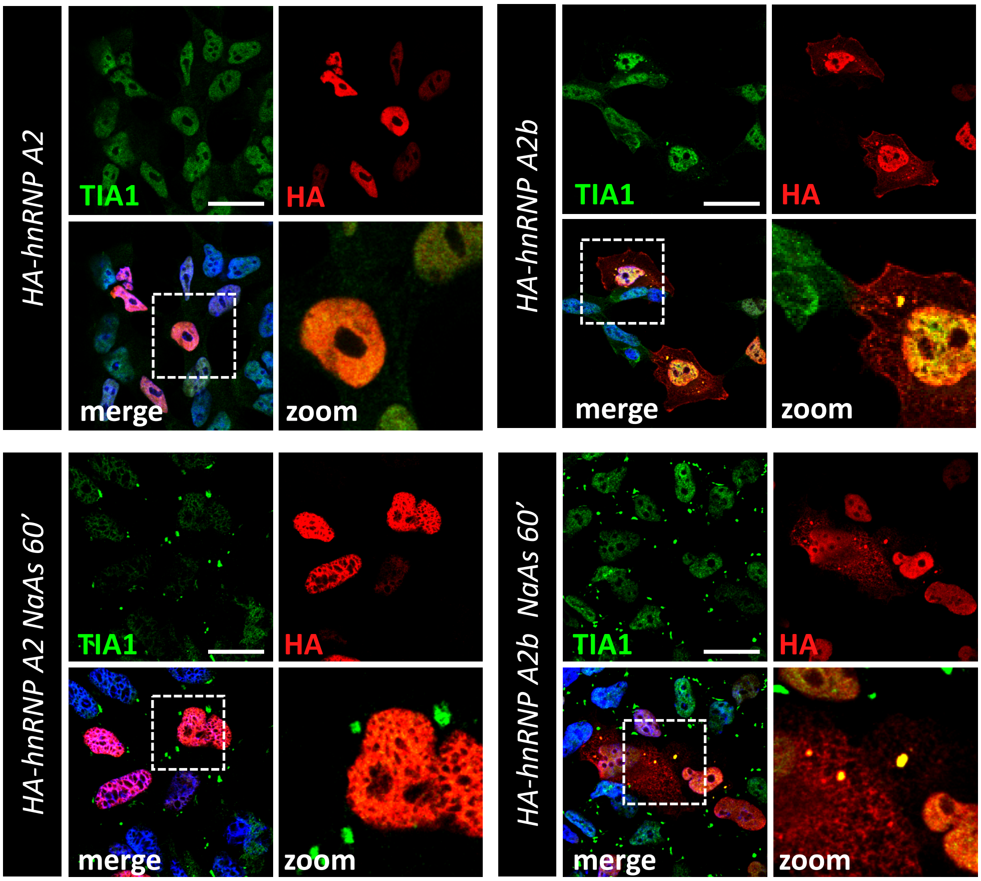

Supplementary FIGURE 5

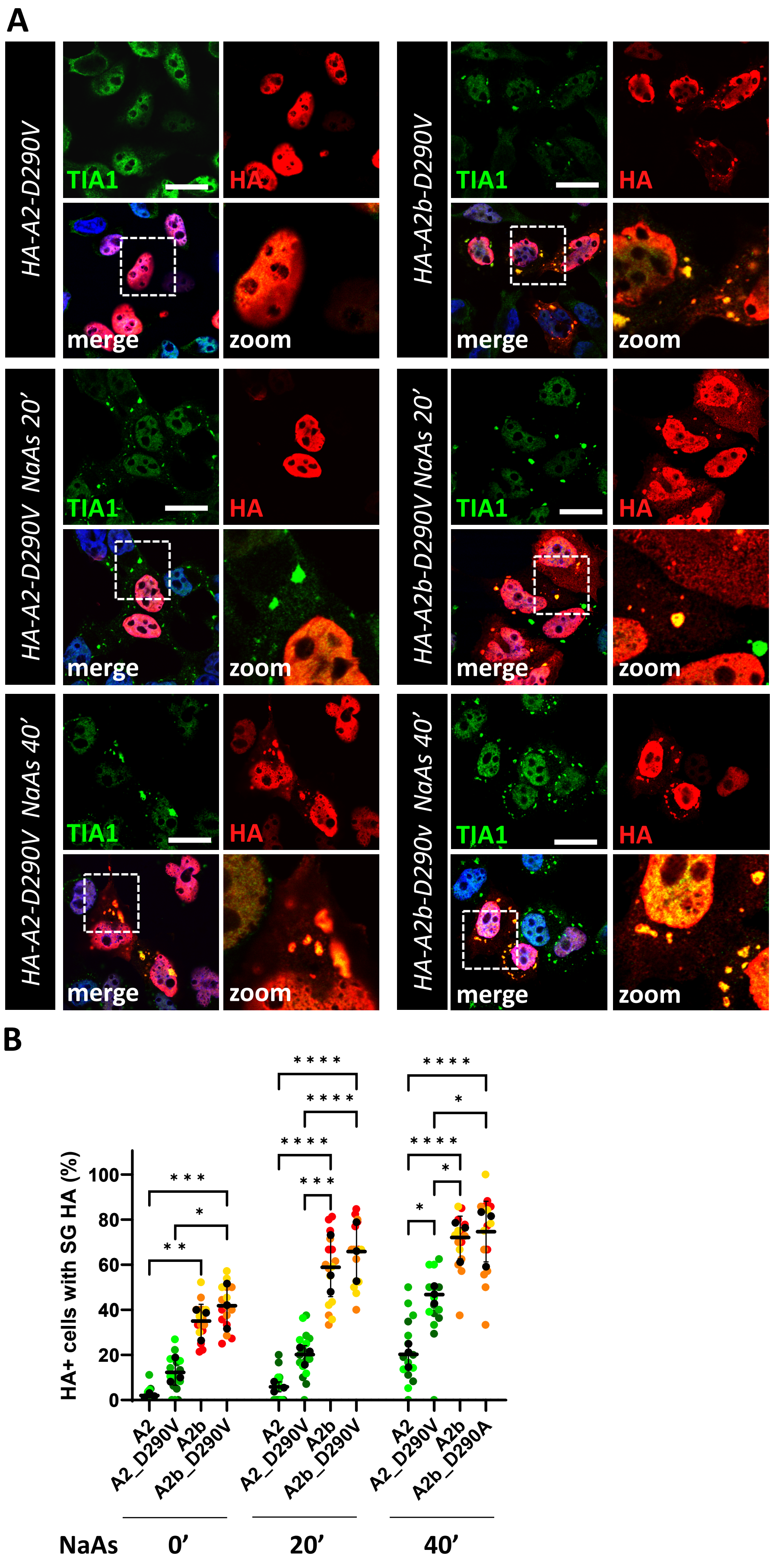

### Supplementary FIGURE 6

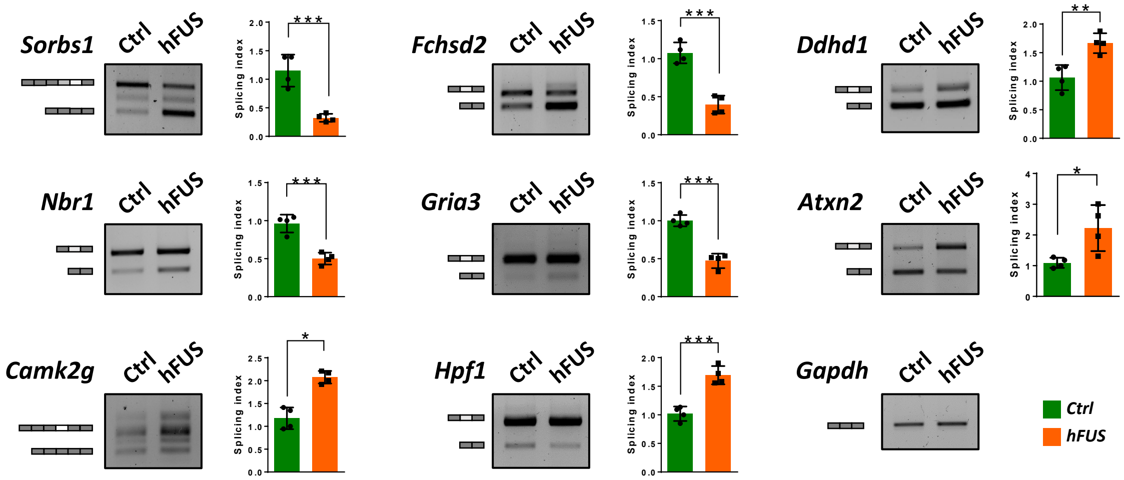

### Supplementary FIGURE 7

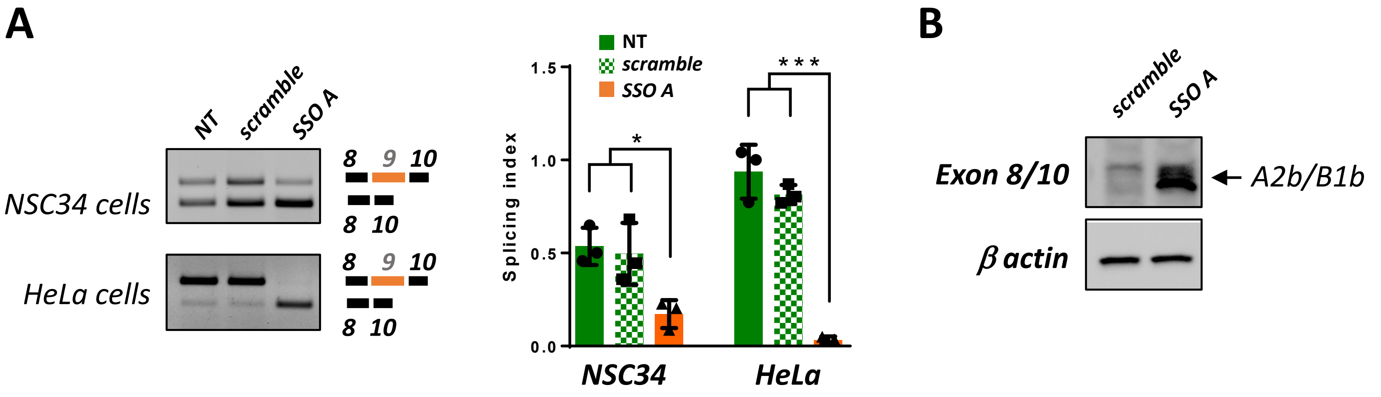

Supplementary FIGURE 8

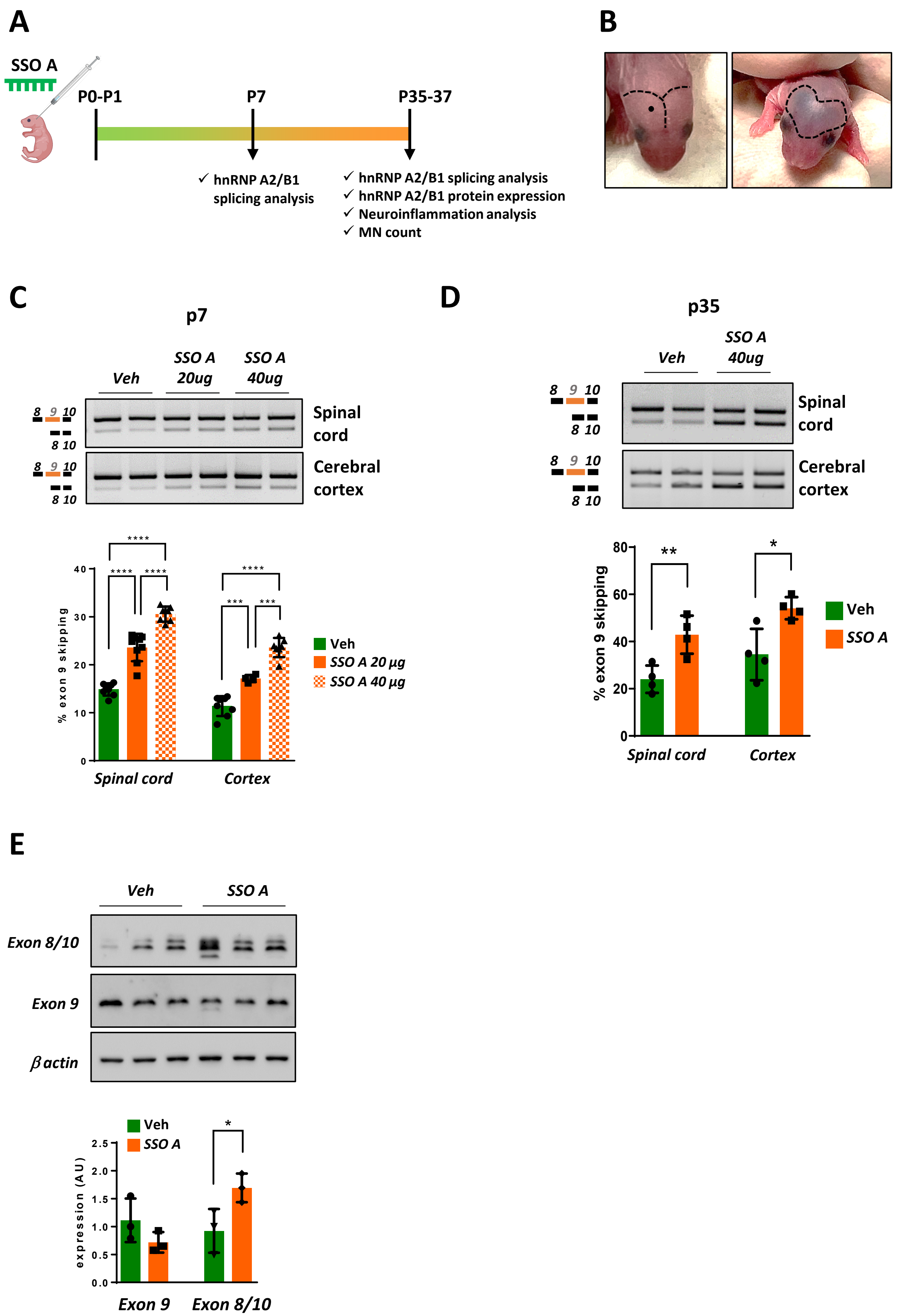

Supplementary FIGURE 9

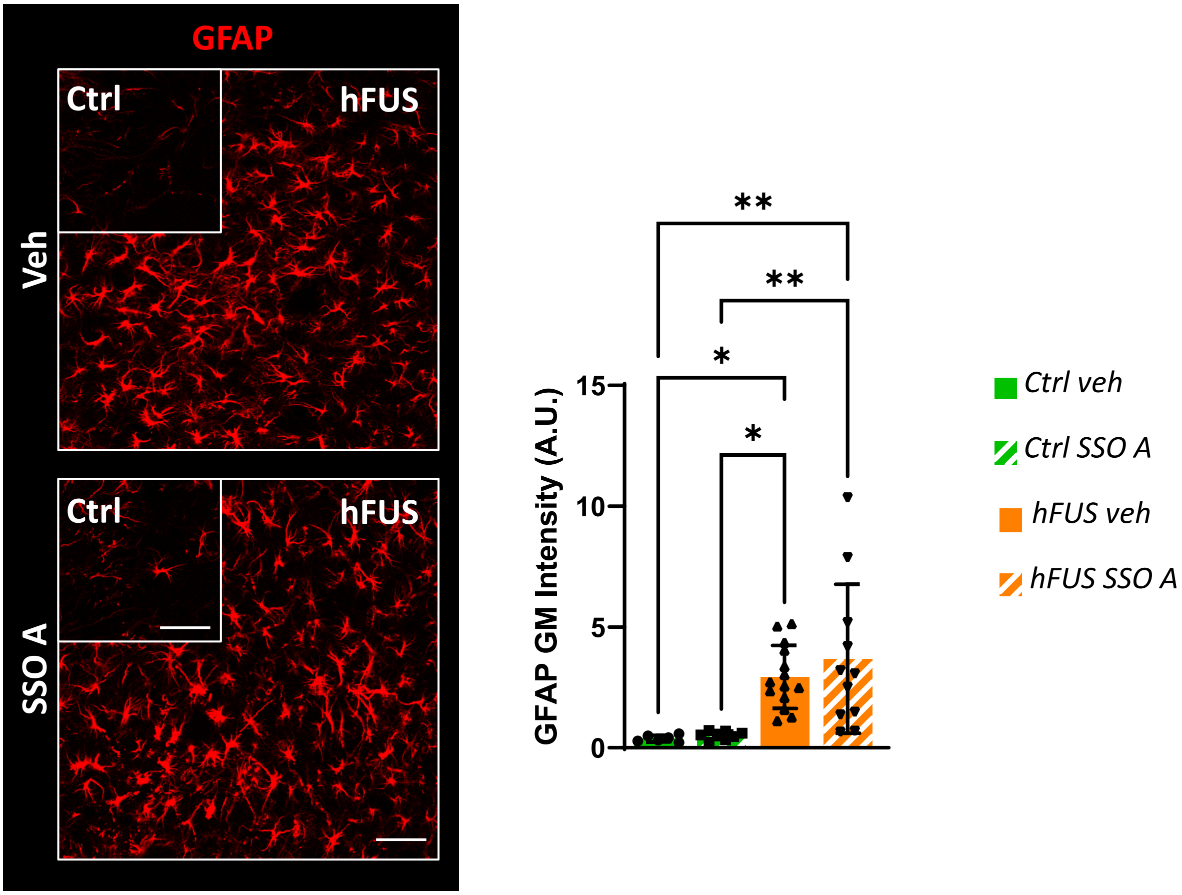

### Supplementary Figure legends

**Supplementary Figure 1. Isoform-specific antibodies recognize distinct hnRNP A2/B1 isoforms.** (A) Schematic representation of the four different hnRNP A2/B1 isoforms generated by the alternative splicing of exons 2 and 9. Black lines represent the epitopes that can be recognized by the isoform-specific antibodies used in this study: anti-exon 9 antibody binds B1 and A2 isoforms; anti-exon 8/10 recognizes B1b and A2b; anti-exon 2 identifies B1 and B1b. (B,C) Protein extracts from HeLa cells transfected with a mock plasmid or plasmids coding for individual hnRNP A2/B1 isoforms were analysed by western blot with the indicated antibodies. Two different preparations of anti-exon 8/10 were tested. Red arrowheads indicate endogenous isoforms.

**Supplementary Figure 2. hnRNP A2/B1 isoforms lacking exon 9 show cytoplasmic localization in primary cortical neurons.** Representative immunofluorescence analysis of primary cortical neurons derived from the cerebral cortex of non-transgenic mice transfected with the HA-tagged hnRNP A2/B1 isoforms and immunolabeled with anti-HA antibody (red) and anti-β-III tubulin antibody (green). Nuclei were detected by DAPI staining (blue). Scale bar: 20 µm.

**Supplementary Figure 3. B1b shows increased localization in stress granules compared to B1 under NaAs treatment.** HeLa cells were transfected with the HA-tagged hnRNP B1 and B1b isoform constructs for 24h, untreated or treated with 0.5 mM sodium arsenite (NaAs) for 20, 40 and 60 minutes. (A) Immunofluorescence analysis of cells untreated or treated with 0.5 mM sodium arsenite (NaAs) for 60 minutes, using an anti-HA antibody (red) and anti-TIA1 antibody (green). Nuclei were detected by DAPI staining (blue). Magnifications of the highlighted areas are also shown (zoom). Scale bar: 20 µm. (B) SuperPlots showing the percentage of cells with the HA signal colocalizing with TIA1-positive stress granules. The distribution of measures from n=3 independent experiments is reported, with each biological replicate color-coded: the mean value from each of the three replicates is represented by black dots, and the mean ± SD of the three replicates is shown as a black line. Statistical significance was calculated by One-way ANOVA test, **p<0.01, ***p<0.001 ****p<0.0001, and the significant differences between B1 and B1b isoforms at the same time point are shown.

**Supplementary Figure 4. A2b shows increased localization in stress granules compared to A2 under NaAs treatment in SH-SY5Y cell line**. SH-SY5Y cells were transfected with the HA-tagged hnRNP A2 and A2b isoform constructs, and after 24h were left untreated or treated with 0.5 mM NaAs for 60' and analysed by immunofluorescence, using an anti-HA antibody (red) and anti-TIA1 antibody (green). Nuclei were detected by DAPI staining (blue). Magnifications of the highlighted areas are also shown (zoom). Scale bar: 20 µm.

**Supplementary Figure 5. Lack of exon 9 enhances the localization of the A2-D290V ALS mutant into stress granules.** (A) HeLa cells were transfected with HA-tagged hnRNP A2-D290V and A2b-D290V constructs and left untreated or treated with 0.5 mM sodium arsenite (NaAs) for 20 or 40 minutes. The cells were analyzed by immunofluorescence using an anti-HA antibody (red) and anti-TIA1 antibody (green). Nuclei were detected by DAPI staining (blue). Magnifications of the highlighted areas are also shown (zoom). Scale bar: 20 µm. (B) SuperPlots showing the percentage of cells with the HA signal colocalizing with TIA1-positive stress granules. The distribution of measures from n=3 independent experiments is reported, with each biological replicate color-coded: the mean value from each of the three replicates is represented by black dots, and the mean ± SD of the three replicates is shown as a black line. Statistical significance was calculated by One-way ANOVA test, and the significant differences between groups at the same time point are shown. *p<0.05, **p<0.01, ***p<0.001, ****p<0.0001.

**Supplementary Figure 6. Alternative splicing of hnRNP A2/B1 target genes is affected in symptomatic hFUS mice.** The alternative splicing of the indicated hnRNP A2/B1 and FUS target genes was assessed through semiquantitative RT-PCR analysis in the spinal cords of symptomatic hFUS mice, along with age-matched non-transgenic control (Ctrl) animals. Representations of constitutive exons (dark grey rectangles) and alternatively spliced exons (light grey and white rectangles) analysed are shown. Bands were quantified by densitometric analysis, and a splicing index was calculated based on the ratio between the upper and the lower band and plotted considering the corresponding ratio in Ctrl mice equal to 1. Data are expressed as means ± SD (n=4 mice/group). Statistical significance was calculated by Student's t-test, *$p<0.05$, **$p<0.01$, ***$p<0.001$.

**Supplementary Figure 7. An SSO inducing exon 9 skipping promotes the accumulation of hnRNP A2b and B1b protein isoforms.** (A) NSC34 and HeLa cells were left untransfected (NT) or transfected with a non-targeting SSO (scramble) or a SSO targeting 5' splice site of alternatively spliced exon 9 of hnRNP A2/B1 (SSO A) at the concentration of 0.5 µM. After 24 hours (HeLa) or 48 hours (NSC34), the alternative splicing pattern of hnRNP A2/B1 exon 9 was analysed by semi-quantitative RT-PCR. Bands were quantified through densitometric analysis, and a splicing index was calculated as the ratio between the upper and lower band. Data are expressed as means ± SD (n=3 independent experiment). Statistical significance was calculated by One-way ANOVA test, *$p<0.05$, ***$p<0.001$. (B) Protein extracts from HeLa cells transfected with control SSO (scramble) or SSO A were analysed after 48 hours with the antibody specifically recognizing the isoforms lacking exon 9 (anti-exon 8/10). The expression of β-actin was measured to normalize protein loading. The migration of the hnRNP A2b/B1b isoforms is indicated by an arrow.

**Supplementary Figure 8. SSO A promotes exon9 skipping *in vivo*.** (A) Schematic illustration of SSO A treatment in vivo. hFUS mice received a single intracerebroventricular (ICV) injection at P0-P1 with 20 µg or 40 µg of SSO A or PBS. Alternative splicing of hnRNP A2/B1 was subsequently analysed in spinal cord and cerebral cortex tissues at 7 (P7) and 35 (P35) days post-ICV injection. (B) Illustration of ICV injection technique. The injection site into one of the lateral ventricles is carefully determined at the midpoint between the eye and the lambda intersection of the skull. The solution rapidly disseminates, achieving dispersion throughout both lateral ventricles within a few minutes post-injection. (C, D) RNAs extracted from spinal cord or cerebral cortex of treated mice with SSO A or PBS (vehicle, Veh) were analysed by RT-PCR to monitor the splicing of hnRNP A2/B1 exon 9. Mice were sacrificed 7 days (C) or 35 days (D) after ICV injection. Bands were quantified through densitometry analysis, and the levels of expression of hnRNP A2/B1 isoforms lacking exon 9 were indicated as the percentage of the total amount of the isoforms containing and excluding exon 9. Data are expressed as means ± SD. At least n=4 mice per group were analysed. Statistical significance was calculated by One-way ANOVA, ***p<0.001, ****p<0.0001. (E) Lumbar spinal cord lysates from hFUS mice treated with PBS (Veh) or 40 μg SSO A were extracted 35 days after ICV injection and subjected to western blot analysis using anti-exon 9 and anti-exon 8/10 antibodies. β-actin was used as a loading control. Data are expressed as mean ± SD considering the relative expression of a Ctrl mice equal to 1.  n=3 mice per group were analysed. Statistical significance was calculated by student's t-test, *p<0.05.

**Supplementary Figure 9. SSO A does not affect astrocytosis.** Spinal cord sections from non-transgenic (Ctrl) and symptomatic hFUS mice treated with PBS (Veh) or 40 μg SSO A were analyzed 35 days after ICV injection and subjected to immunofluorescence staining with an antibody against GFAP (red). N = 3/4 animals for group, at least four sections for animal, were analysed. Statistical significance was calculated using ANOVA. Scale bar: 50 μm, insert scale bar: 4 μm.

**Supplementary Table 1. Sequences of primers used in semi-quantitative RT-PCR.**

| Gene | Primers |
|---|---|
| hnRNP A2/B1 (E1-E3)<br>(Exon 2 splicing assay) | Fwr: 5′ - TCTTGGCCATCGCCTGCT - 3′<br>Rev: 5′ - TCCCCATTGCTCATAGTAGT - 3′ |
| hnRNP A2/B1 (E7-E10)<br>(Exon 9 splicing assay) | Fwr: 5′ - GGATTCTCGTGGTGGCGG - 3′<br>Rev: 5′ - CATATGGTCCTCCCATGTTC -3′ |
| Sorbs1 (E14-E19 | Fwr: 5′ - ATGATTCTGATGTCCATTCCC -3′<br>Rev: 5′- CTTTCTCTAAATCTGCCTCTAAC- 3′ |
| Nbr1 (E16-E18) | Fwr: 5′ - TCCTTTGAGCTGCTGGATATA -3′<br>Rev: 5′ - AGGGTT GTG GTC TGG AGCA - 3′ |
| Camk2g (E12-E17) | Fwr: 5′ - GGTGCCATCCTCACAACCAT - 3′<br>Rev: 5′- CTTCGTCTTCTGTAGTGGTG - 3′ |
| Tbp (E5-E7) | Fwr: 5′ - ATCATGAGAATAAGAGAGCC - 3′<br>Rev: 5′ - CTGGGTTTGATCATTCTGTAG - 3′ |
| Fchsd2 (E10-E12) | Fwr: 5′ - CGGGACTATAACCTTCAGCTG - 3′<br>Rev: 5′ - TGTGCTCACGTGCCACACGT - 3′ |
| Gria3 (E13-E15) | Fwr: 5′ - GGAAGTCCAAGGGAAAGTTC - 3′<br>Rev: 3′ - TGCCACATTGCTCAGGCTTA - 3′ |
| Hpf1 (E2-E4) | Fwr: 5′ - TCTGAGGCTGATGTCTCCAGT - 3′<br>Rev: 5′ - GCAGCAAACACATTGTCTCCA - 3′ |
| Ddhd1 (E11-E13) | Fwr: 5′ - CAGAGTAAAGATGATAGCCTC - 3′<br>Rev: 5′ - TCAGAGTGAACCTAGGCTG - 3′ |
| Sp100 (E10-E13) | Fwr: 5′ - GCATATTCCAGAAAACCTCC - 3′<br>Rev: 5′ - GATTGGCACTTCTTGCTAGG - 3′ |
| Atxn2 (E9-E11) | Fwr: 5′ - TCAAGAGCTGCTTCTCACA - 3′<br>Rev: 5′ - AGGAGCAGCTGCTTCAC - 3′ |
| eIF4h (E4-E6) | Fwr: 5′ - TTGCAGAAGGCAGAAAACAAG - 3′<br>Rev: 5′ - TCTGAAGTCCATGTTGGAGCC - 3′ |
| GAPDH | Fwr: 5′ - ACCACCATGGAGAAGGCCGGG - 3′<br>Rev: 5′ - CAGTGATGGCATGGACTGTGG - 3′ |

**Supplementary Table 2. List of overlapping genes whose alternative splicing is misregulated upon FUS depletion (FUS ASO), FUS knockout (FUS KO) and hnRNP A2/B1 depletion (A2/B1 ASO).**

| | **A2B1 ASO, FUS ASO and FUS KO** | **A2B1 ASO and FUS KO** | **A2B1 ASO and FUS ASO** |
|---|---|---|---|
| 1 | Fchsd2 | Ep400 | Hmgn3 |
| 2 | Tbp | Smc6 | Rbm3 |
| 3 | Ogdh | Atxn2 | Stau2 |
| 4 | Nbr1 | Ipo11 | Sytl2 |
| 5 | Ptprz1 | Camk2g | Fus |
| 6 | 2700029M09Rik (Hpf1) | Sp100 | Ezh2 |
| 7 | Eif4h | Arnt | Fgfr3 |
| 8 | Clk4 | Carf | Kcnip4 |
| 9 | Sorbs1 | | Bcas1 |
| 10 | Gria3 | | Eno2 |
| 11 | Ddhd1 | | Mboat7 |
| 12 | Ttc3 | | Abcc5 |
| 13 | Pax6 | | Eif4g2 |
| 14 | D4Wsu53e | | 5930434B04Rik |
| 15 | | | R3hdm1 |
| 16 | | | Wsb1 |
| 17 | | | Morc3 |
| 18 | | | Pex5l |
| 19 | | | Enox2 |
| 20 | | | Ttc14 |
| 21 | | | Tia1 |
| 22 | | | Phc3 |
| 23 | | | Ppap2c |
| 24 | | | Etl4 |
| 25 | | | Clk1 |
| 26 | | | Plch2 |
| 27 | | | Chpt1 |
| 28 | | | Cd68 |
| 29 | | | Macrod2 |
| 30 | | | Tpd52l2 |
